# Controlling gene activation by enhancers through a drug-inducible topological insulator

**DOI:** 10.1101/534073

**Authors:** Taro Tsujimura, Osamu Takase, Masahiro Yoshikawa, Etsuko Sano, Matsuhiko Hayashi, Kazuto Hoshi, Tsuyoshi Takato, Atsushi Toyoda, Hideyuki Okano, Keiichi Hishikawa

**Author notes:** **Corresponding authors** Taro Tsujimura, Keiichi Hishikawa.

## Abstract

While regulation of gene-enhancer interaction is better understood, its application remains limited. Here, we reconstituted arrays of CTCF binding sites and devised a <u>s</u>ynthetic <u>t</u>opological insulator with <u>t</u>etO for <u>ch</u>romatin-engineering (STITCH). By coupling STITCH with tetR linked to the KRAB domain to induce heterochromatin and disable the insulation, we developed a drug-inducible system to control gene activation by enhancers. We applied this to dissect *MYC* regulation in human pluripotent stem cells. Insertion of STITCH between *MYC* and the enhancer down-regulated *MYC* and affected its target transcriptome. Progressive mutagenesis of STITCH led to preferential escalation of the gene-enhancer interaction, corroborating the strong insulation ability of STITCH. The STITCH insertion altered epigenetic states around *MYC*. Time-course analysis by drug induction uncovered deposition and removal of H3K27me3 repressive marks follows and reflects, but does not precede and determine, the expression change. Thus the tool provided important insights in gene regulation, demonstrating its potency.

## Introduction

Interaction of genes and enhancers is greatly affected by architectural proteins that bind to chromatin and organize folding of the genome (Dekker et al., 2017). Most notably, CTCF mediates loop formation of chromatin as an anchor of a cohesin complex, which physically bundles two distant loci of the genomic DNA (Kagey et al., 2010; Phillips-Cremins et al., 2013; Wendt et al., 2008). The genome wide contact maps of chromatin show that the CTCF binding sites often demarcates boundaries of so called contact domains or topologically associating domains, where chromatin association takes place more preferentially inside than outside (Dixon et al., 2012; Phillips-Cremins et al., 2013; Rao et al., 2014). The looping between two CTCF binding sites are established exclusively when the two sites are in the converging orientations with each other (Rao et al., 2014). Loss of cohesin or CTCF resulted in disappearance of contact domains (Nora et al., 2017; Rao et al., 2017; Schwarzer et al., 2017). According to the extrusion model, the cohesin ring extrudes the chromatin fiber from a loaded site and pauses at a CTCF binding site that is oriented towards the ring (Fudenberg et al., 2016; Sanborn et al., 2015). This model is widely accepted as the underlying mechanism for the formation of the loops and contact domains. On the other hand, several studies have shown that the CTCF boundaries limit the action ranges of enhancers and thus restrict the enhancer targets to genes within the same contact domains as the enhancers reside in (Dowen et al., 2014; Lupiáñez et al., 2015; Symmons et al., 2014; Tsujimura et al., 2015; 2018). Thus CTCF demarcates contact domains and interferes with gene-enhancer interaction when located in between, though the mechanism of the interference of the interaction is not entirely clear.

While the relationship between CTCF binding, chromatin conformation, and gene-enhancer interaction has become evident, its application to control gene expression is limited. Forced bridging between enhancers and genes via DNA recognizing domains such as zinc-finger proteins and CRISPR/dCas9 have been successful to induce gene expression (Deng et al., 2014; Morgan et al., 2017). However, importing these systems to a given locus is not easy, as prior characterization of enhancers and careful design of DNA recognizing domains is essential for each locus. In this sense, using CTCF binding sites as an enhancer blocker is a very appealing approach, as it can blunt gene activation as long as the element is inserted between the gene and the enhancers without knowing precisely where the enhancers are. The insulator element identified in the chicken *β-globin* locus, which harbors a CTCF binding site (Bell et al., 1999), has been utilized in heterologous systems (Bessa et al., 2014). From our view, however, those elements have not been spread widely as tools to control gene expression, probably because the underlying mechanism was not fully understood and convincing explanation for its general utility was missing when they were reported. However, engineering the genome based on the CTCF function has possibility to add a new layer to the techniques of artificially controlling gene expression, and should open a new avenue in diverse fields of biology and medicine.

The *Tfap2c-Bmp7* locus in mice is partitioned into two contact domains by a region termed TZ (Tsujimura et al., 2015; 2018). The TZ limits target ranges of enhancers at the locus (Tsujimura et al., 2015). Two arrays of CTCF binding sites in divergent configuration composing the TZ provide the strong ability to block chromatin contacts (Tsujimura et al., 2018). Taking advantage of the well-characterized nature of the TZ, in this study, we developed a new system to control interaction between a gene and an enhancer. We first reconstituted the CTCF binding sites of the TZ as a short DNA cassette, which successfully functioned as an enhancer blocker heterologously at the *MYC* locus in human induced pluripotent stem cells (iPSCs). Further we added a feature that enables epigenetically controlling the blocking activity of the cassette in a drug-inducible manner. Here we describe the system, demonstrate its utility to study gene regulation by enhancers, and discuss the future potential of the system.

## Results

### STITCH blocks the interaction of *MYC* with the enhancer when inserted in between

The TZ consists of seven binding sites of CTCF: they are L1, L2, L3, L4, R1, R2 and R3, arrayed in this order from the *Tfap2c* side to the *Bmp7* side (Figure 1A) (Tsujimura et al., 2018). L1-L4 are oriented towards *Tfap2c* and collectively referred to as L, while R1-R3 are towards *Bmp7* and referred to as R. Though the seven sites are constantly called as peaks of CTCF binding in different cell types by ChIP-seq (Chromatin immunoprecipitation followed by sequencing) with cross-linking, native-ChIP (nChIP) failed to detect CTCF binding at L1 and L4, suggesting the binding there is weak or indirect (Tsujimura et al., 2018). We extracted the 178- or 179-bp DNA sequences including the consensus motif sequence for CTCF binding and concatenated them as a short DNA cassette. We embedded the core sequence of the tetracycline operator (tetO) at four different positions within the cassette. tetO is bound by the tetracycline repressor (tetR), but not in presence of doxycycline (DOX), and thus allows recruitment of an effector protein to the cassette in a drug-dependent manner (Gossen and Bujard, 1992). We also put a puromycin-resistant gene sandwiched by two loxP sites for the sake of efficient targeting (Figure 1A, Table S2). We expect that the CTCF binding sites of the cassette would recruit CTCF and function as a topological insulator, and that the tetO/tetR system would enable epigenetically modifying the insulation activity. We named the cassette as <u>S</u>ynthetic <u>T</u>opological Insulator with <u>T</u>etO for <u>Ch</u>romatin-engineering (STITCH) (Figure 1A).

**Figure 1.**
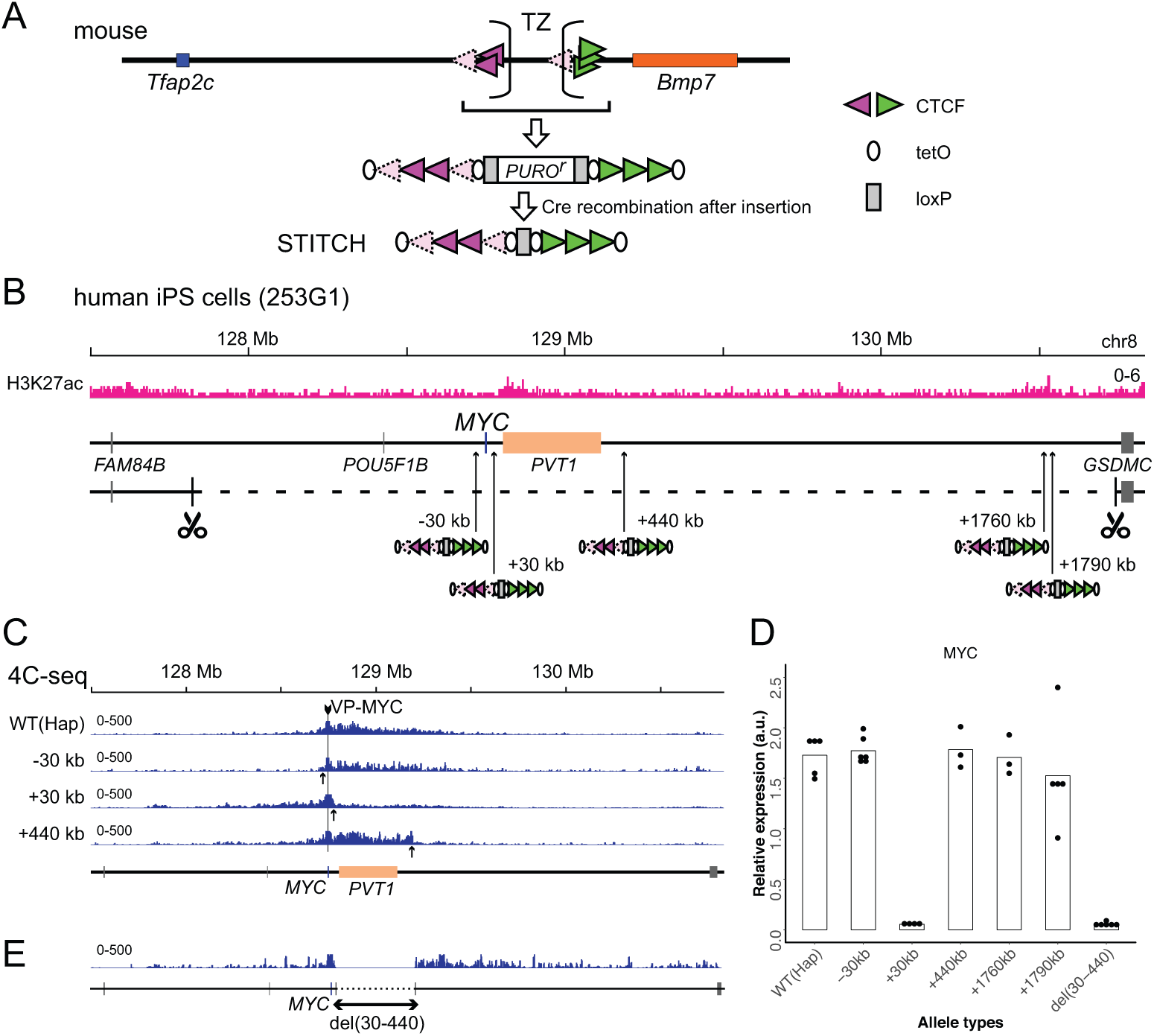
Serial insertion of STITCH around *MYC* localized the enhancer. (A) Design of STITCH and scheme of inserting the cassette. After recombination of the two loxP sites (rectangles), the puromycin resistant gene is removed. Directionality of the CTCF binding motifs is represented by the orientation and color of the triangles. Note that binding of CTCF at L1 and L4 is not detected by nChIP, as represented by the paled color. The ovals represent tetO. (B) The H3K27ac profile and the insertion sites of STITCH around *MYC* in the human iPS cells. The 3-Mb region deleted from one of the two alleles to make “Hap” is indicated by the dashed line, flanked by scissors that indicate the target sites of CRISPR/Cas9. The numbers in the insertion names indicate the distance from *MYC*. (C) The 4C-seq profiles from VP-MYC2 of the wild type (Hap) and STITCH-30kb, +30kb and +440kb alleles. (D) Relative *MYC* expression levels normalized with *ACTB* expression in the different alleles. Each dot represents replicate clones (see Materials and Methods for details). The bars represent their means. (E) The 4C-seq profile of del(30-440) from VP-MYC2.

*MYC* is highly expressed in human pluripotent stem cells (Knoepfler, 2008). As *MYC* expression is regulated by long-range enhancers in various cell types, we thought that *MYC* expression in the stem cells should be also dependent on long-range enhancers (Bahr et al., 2018; Cho et al., 2018; Dave et al., 2017; Herranz et al., 2014; Hnisz et al., 2013; Lovén et al., 2013; Pulikkan et al., 2018; Shi et al., 2013; Sur et al., 2012; Uslu et al., 2014; Zhang et al., 2016). Hence we used the iPSC line 253G1, which was generated via retroviral transduction of *OCT4*, *KLF4* and *SOX2* but without *MYC*, to test the functionality of STITCH (Nakagawa et al., 2007). Since the diploidy would hamper the following genome editing procedures, we first deleted one allele of the 3-Mb region around *MYC* as described before to make the locus locally haploid, and termed the clone as “Hap” (Figure 1B) (Tsujimura et al., 2018). Then we inserted STITCH into five different positions of the remaining allele of the locus: “STITCH+30kb”, “STITCH+440kb”, “STITCH+1760kb” and “STITCH+1790kb” have the STITCH insertions away from the *MYC* promoter for the indicated distances to the telomeric side of the q arm of the chromosome (the right side on the map, Figure 1B); “STITCH-30kb”, at the 30-kb upstream from the *MYC* (the left side, Figure 1B). STITCH+30kb and STITCH+440kb flank the neighboring long non-coding RNA (lncRNA) gene *PVT1*. Broad deposition of the enhancer associated histone modification H3K27ac is observed within the *PVT1* region. STITCH+1760kb and STITCH+1790kb flank a peak of H3K27ac (Figure 1B).

We first performed 4C-seq (Circular chromatin conformation capture assay followed by deep sequencing) from the *MYC* promoter as a viewpoint to see how STITCH impacts on the chromatin conformation. We designed two sets of primers around the *MYC* promoter as viewpoints of 4C-seq (VP-MYC1 and VP-MYC-2, see Figure S1A). In the wild type allele of Hap, *MYC* mainly contacts with the *PVT1* region and around (Figure 1C and Figure S1B). In STITCH+30kb, STITCH+440kb and STITCH-30kb, the contacts were blocked at the inserted positions of STITCH as expected (Figure 1C and Figure S1B). We then extracted RNA from the cells and measured the *MYC* expression levels with quantitative PCR (qPCR). We found that only STITCH+30kb strongly down-regulated the *MYC* expression, while the others did not (Figure 1D). These results suggest that the region between STITCH+30kb and STITCH+440kb (+(30-440)kb region) possesses the enhancer activity for the *MYC* expression. We made a deletion clone of the region, termed del(30-440) (Figure 1E and Figure S2A). While the 4C contact extended further away from the deleted region (Figure 1E and Figure S1B), *MYC* was strongly down-regulated by the deletion (Figure 1D), showing the region contains the enhancer. nChIP-seq in STITCH+30kb confirmed that each of the biding sites of STITCH, except L1 and L4, was bound by CTCF (Figure S1C). Thus STITCH recruits CTCF and blocks interaction between a gene and an enhancer when located in between as an insulator.

### Insulation and deletion of the enhancer resulted in similar transcriptome profiles

We employed RNA-seq to see how the insulation (STITCH+30kb) and deletion (del(30-440)) of the enhancer affects the transcriptome of the cells (Figure 2 and Figure S3). We prepared libraries from three replicates for each configuration. Consistently with the qPCR assay (Figure 1D), strong down-regulation of *MYC* was confirmed in both STITCH+30kb and del(30-440) (Figure 2A and Figure S3). *PVT1* expression was not altered in STITCH+30kb (Figure 2A and Figure S3). We did not observe other detectable expression changes around the *MYC* locus in either STITCH+30kb or del(30-440) (Figure 2A). Comparison of the transcriptome with DESeq2 (Love et al., 2014) detected 564 and 222 down-regulated genes and 839 and 350 up-regulated genes in STITCH+30kb and del(30-440), respectively (Figure 2B-F). Among those, large fractions (142 and 215 genes, for down- and up-regulation, respectively) were common between the two alleles (Figure 2E and F). Notably, the commonly down-regulated genes are highly enriched with genes encoding regulators of ribosome assembly and translation as well as those involved in metabolism particularly for cholesterol, which are known target groups of *MYC* in various systems (Figure 2G) (Hofmann et al., 2015; Uslu et al., 2014; van Riggelen et al., 2010; Zeller et al., 2006) We found several GO terms that are related to specialized cell functions as enriched in commonly up-regulated genes, but the enrichment was not that high (Figure 2H). Importantly, the transcriptome comparison between STITCH+30kb and del(30-440) called much less number (261) of differentially expressed genes, showing their difference is limited (Figure 2D). Thus our results show that both the insulation and deletion of the *MYC* enhancer similarly down-regulated *MYC* and affected its downstream targets.

**Figure 2.**
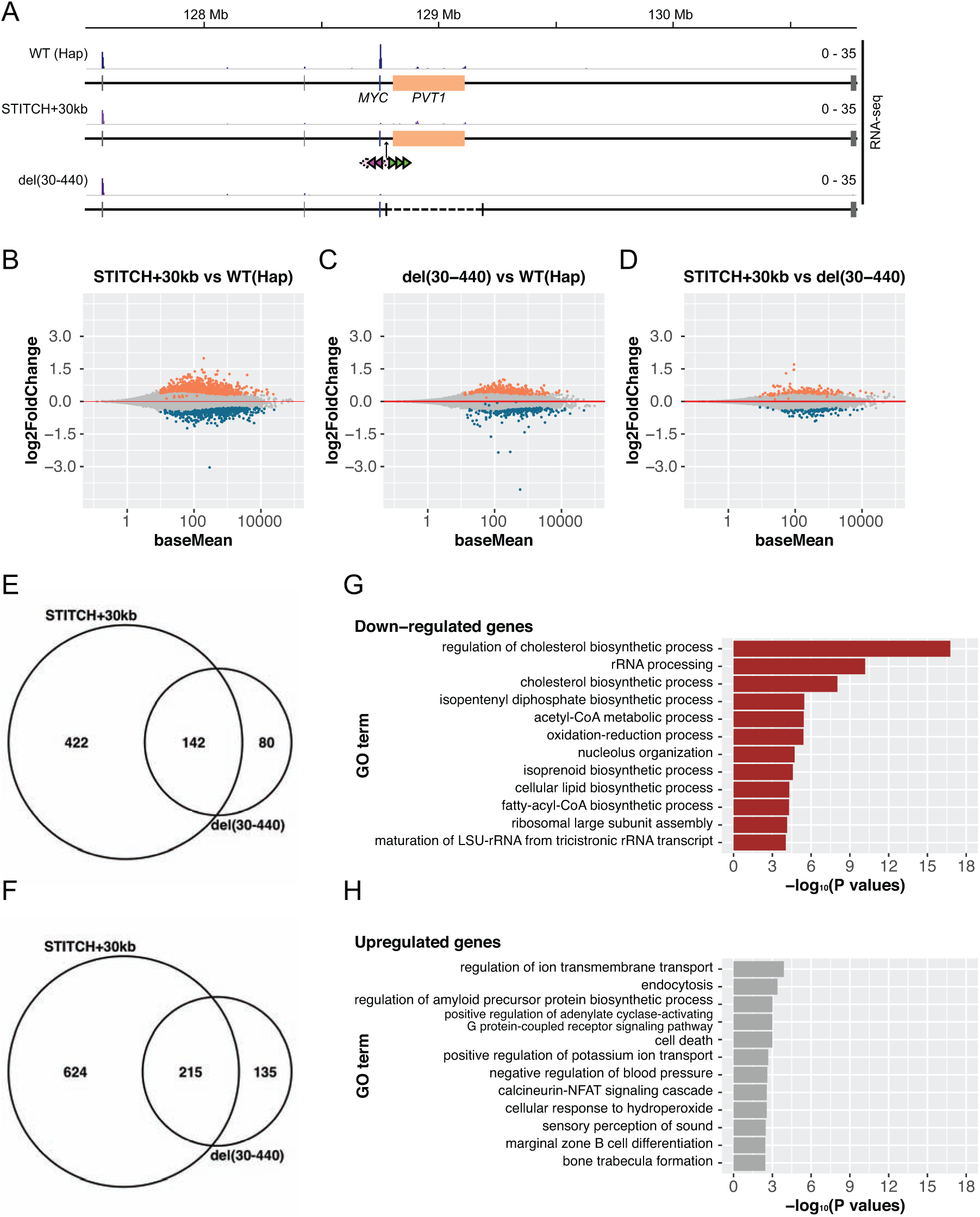
Transcriptome analysis of Hap, STITCH+30kb and del(30-440) (A) Tracks of RNA-seq from Hap, STITCH+30kb and del(30-440) around the *MYC* locus. (B-D) MA-plots of RNA-seq to compare STITCH+30kb vs Hap (B), del(30-440) vs Hap (C) and STITCH+30kb vs del(30-440) (D). Differentially expressed genes (FDR < 0.1) are marked by colors (orange for up-regulated genes and dark blue for down-regulated ones). (E and F) Venn diagrams showing overlaps of differentially down-regulated (E) and up-regulated (F) genes in STITCH+30kb and del(30-440). (G and H) GO enrichment analysis of the commonly down-regulated (G) and up-regulated (H) genes in STITCH+30kb and del(30-440).

### Titrating blocking activity of STITCH by serial mutations of the CTCF binding sites

Deletion and inversion of the CTCF binding arrays impaired the blocking activity of the TZ at the endogenous locus in the mouse embryonic stem (ES) cells, showing that the diverging configuration most strongly blocks contacts (Tsujimura et al., 2018). In the present study, to test if this holds true in the reconstituted cassette, we made deletion of each CTCF array, L (delL) and R (delR), inversion of R (invR), deletion of the middle five binding sites from L2 to R2 (del(L2-R2)), and deletion of the six sites but for R3 (del(L1-R2)) in STITCH+30kb. We also obtained deletion and inversion of the whole of STITCH (del(L1-R3) and inv(L1-R3)) (Figure 3A and Figure S2B-E).

**Figure 3.**
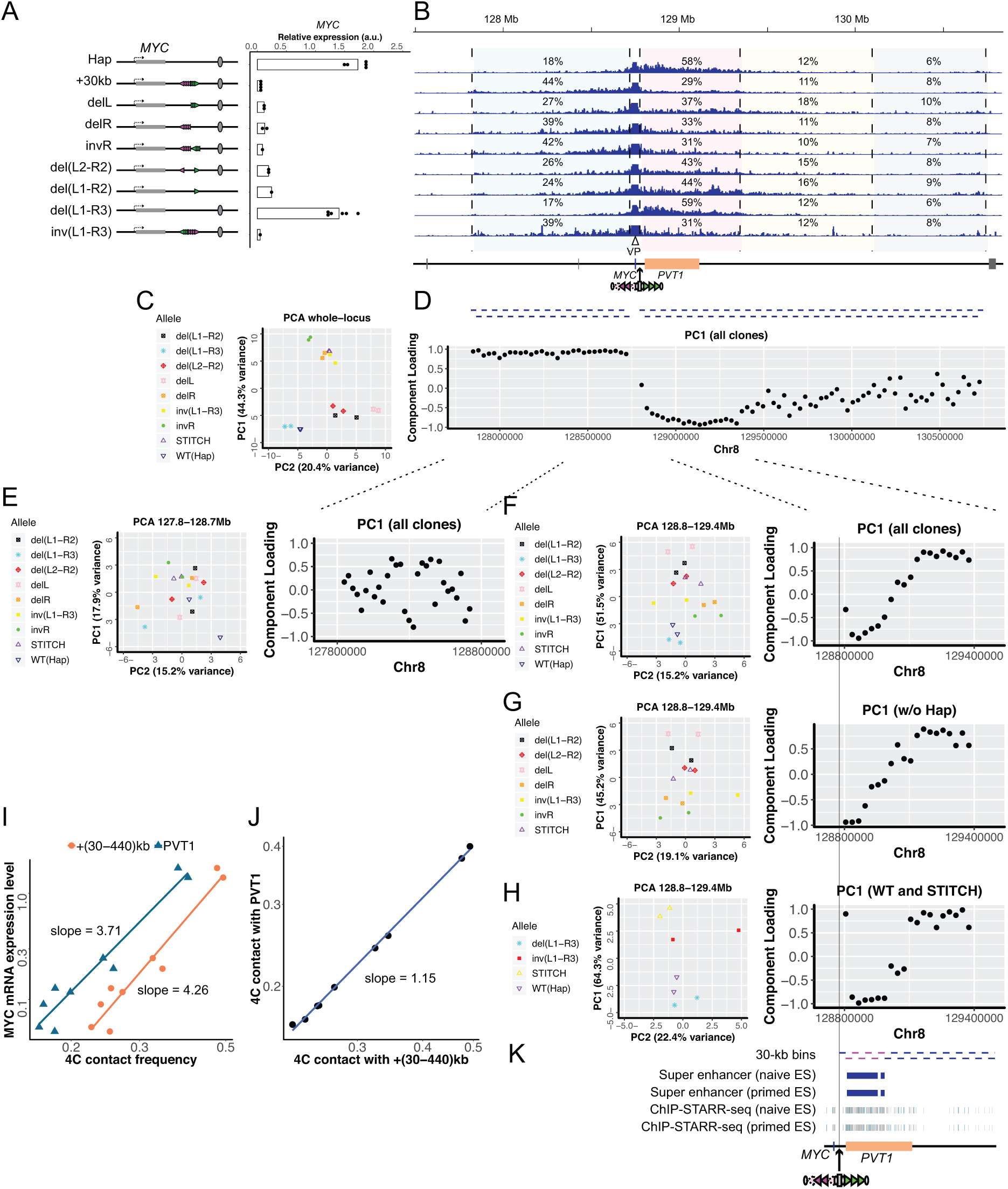
*MYC* expression and 4C-seq profiles in serially mutated STITCH alleles. (A) Configurations of CTCF binding sites of mutated STITCH alleles and a plot showing their *MYC* expression levels. Each dot represents replicate clones (see Materials and Methods for details). Note the data of Hap and STITCH+30kb are same as Figure 1D. Bars indicate means of the replicates. (B) 4C-seq profiles from VP-MYC2 in the different alleles. The numbers are the ratios of the mapped reads to the indicated regions within the 3-Mb region, except for the 10-kb region from the viewpoint. Below the coordinate map, blue bars indicate bins (each 30 kb) for PCA in (C-H) and Figure S5. (C) PCA plot of all the clones using the normalized counts in all the bins of the whole locus. (D) Component loadings of PC1 in the PCA in (C) are plotted along the coordinate for each bin. (E) PCA of all the clones using only the bins of the left 900-kb regions (left) and the PC1 component loading plot (right). (F-H) PCA with all the clones (F), with clones but for the non-blocking alleles (Hap and del(L1-R3)) (G), and only with the non-blocking alleles, the original STITCH and inv(L1-R3) using the bins of the right 600-kb region (H) (left), and the corresponding PC1 component loading plots (F-H) (right). (I) A log-log plot of the *MYC* expression levels against the 4C contact frequencies in the +(30-440)kb region (orange) and the *PVT1* region (dark blue) for each clone. Note the difference of the slopes. (J) A log-log plot of the 4C contact frequencies in the *PVT1* region against the +(30-440)kb region. (K) Tracks of the super-enhancers and ChIP-STARR-seq plots reported in (Barakat et al., 2018) along with the 30-kb bins of the right 600-kb region. The six bins with lowest values of component loadings in (H) are depicted with pink.

The *MYC* expression levels in delL and delR were slightly increased from the original STITCH allele (Figure 3A). invR also increased it but to a lesser extent (Figure 3A). del(L2-R3) and del(L1-R3) up-regulated the expression even more, but much less than the wild type Hap allele (Figure 3A). The *MYC* expression in del(L1-R3) was comparable to that of Hap, showing that the gene activation could be safely recovered upon removal of the CTCF binding sites (Figure 3A). inv(L1-R3) exhibited same degree of repression as STITCH+30kb, showing that STITCH blocks enhancer activation regardless of the orientation of the insertion as a whole (Figure 3A).

We next examined the 4C contact profiles of the *MYC* promoter in these mutation alleles (Figure 3B-J, Figure S4). The contact frequency with the +(30-440)kb enhancer region was apparently changed depending on the configuration (Figure 3B, Figure S4A). The original STITCH and inv(L1-R3) most strongly reduced the contacts. invR showed slightly more of contacts there, but not as much as delL and delR, indicating first that the divergent configuration is the strongest way to block contacts as in the TZ and secondly that the more CTCF binds there, the more strongly it blocks contacts (Figure 3B, Figure S4A). del(L2-R3) and del(L1-R3) further recovered the contact frequency (Figure 3B). Thus the gene expression level and the contact frequency are well correlated. We noted that the expression level fits with a power-law model with the contact frequency of the +(30-440)kb region with a scaling exponent of 4.1-4.3 (Figure 3I and Figure S4B). The Spearman’s rank correlation coefficients were 0.92 and 0.90 for VP-MYC1 and VP-MYC2, respectively.

The redistribution of the contacts by the mutations largely took place between the left and the right sides from *MYC*/STITCH. However, the contact pattern within the right side also seemed varied (Figure 3B and Figure S4A). To quantitatively understand the difference of the contact profiles among the different alleles, we performed the principal component analysis (PCA) for the 4C contact frequencies of 30-kb bins within a given region (Figure 3B-H, Figure S5). We first analyzed the frequencies within the whole *MYC* locus for all of the alleles above. The PCA plot well segregated the non-blocking alleles (Hap and del(L1-R3)) from the other blocking ones, especially the original STITCH, inv(L1-R3) and those with multiple CTCF binding sites orienting to the left side, i.e. invR and delL, along the PC1 axis (Figure 3C). Plotting the component loadings of each bin along the coordinate shows that the segregation is mostly explained by lower and higher frequencies in the left side region, and higher and lower frequencies in the 570-kb region from the +30kb site to the right side, of the non-blocking and the blocking alleles, respectively (Figure 3D). Also the high score of invR and delR along the PC1 indicates that the folding bias of chromatin due to the multiple CTCF bindings towards the left side also accounts for the segregation (Figure 3C and D).

To uncouple the simple blocking effect and the folding bias by the CTCF directionality, we performed PCA against subsets of the clones (Figure S5A and B). First we removed the Hap/del(L1-R3) clones to reduce the variance due to the blocking effect. Then, the clones with leftward binding sites of CTCF came to the upper side and those with rightward were at the lower side along the PC1 (Figure S5A). The component-loading plot changed so that the adjacent regions to the +30kb site in the right side showed positive values and regions located further broadly showed negative values, indicating that the CTCF orientations at the STITCH largely biases the 4C contact pattern between leftward and rightward (Figure S5A). We next used only the Hap/del(L1-R3), the original STITCH and inv(L1-R3) clones for PCA (Figure S5B). Then again the segregation was observed between the non-blocking and blocking alleles (Figure S5B). The close association between the original STITCH and its inversion on the plot suggests that the bias due to the CTCF orientations is little here (Figure S5B). The strongest components of the PC1 were the whole left side region and the 570-kb region to the right side from the insertion site (Figure S5B). The region beyond the 570-kb (or particularly beyond the +690kb position) did not contribute to the segregation along the axis much (Figure S5B). In fact the contact frequencies are similar between them there (Figure 3B and Figure S4A).

### Enhanced association of *MYC* with the super-enhancer upon removal of STITCH

We further performed PCA for each of the left 900-kb region and the right 600-kb region, respectively (Figure 3E-H, Figure S5C and D). The PCA plot for the left side did not show apparent segregation (Figure 3E, Figure S5C and D). By contrast, PCA for the right 600-kb region showed a peculiar tendency (Figure 3F-H). When the subset clones without the non-blocking Hap/del(L1-R3) alleles were analyzed, they were basically ordered by the directionality and the number of the CTCF binding sites along the PC1 (Figure 5G). The component-loading plot consistently exhibited the bias between the leftward and the rightward directions (Figure 5G). The PCA only with the Hap/del(L1-R3) and the STITCH/inv(L1-R3) clones again showed segregation between the non-blocking and blocking alleles along the PC1 axis (Figure 5H). The pattern of the component loadings was notable. The association with the *PVT1* region, especially with the first 180-kb region, accounts for the Hap/del(L1-R3) clones, and that with the other remaining regions for the STITCH/inv(L1-R3). A recent study detected enhancer activity within the *PVT1* region by ChIP-STARR-seq in human embryonic stem cells, and also called the 180-kb region as a super-enhancer, a large stretch of enhancer-like regions (Figure 3K) (Barakat et al., 2018). These results suggest that *MYC* has preferential contacts with the super-enhancer/*PVT1* region in absence of the CTCF insulation. We in fact found that the power-law scaling of the *MYC* expression with the contact frequency with the *PVT1* region has a scaling exponent of 3.6-3.7, which is slightly less than with the +(30-440)kb region (Figure 3I). Consistently, the contact with *PVT1* scales with that with the +(30-440)kb region, with an exponent factor 1.14-1.15 (Figure 3J). Thus titration of STITCH insulation revealed that the contact of *MYC* with the super-enhancer/*PVT1* region is enhanced when the insulation is absent. Or in other words, the presence of CTCF insulation effectively limits the gene-enhancer contact, which may in part account for the reason why action ranges of enhancers are so strongly limited by the CTCF insulators here and in other contexts.

**Figure 4.**
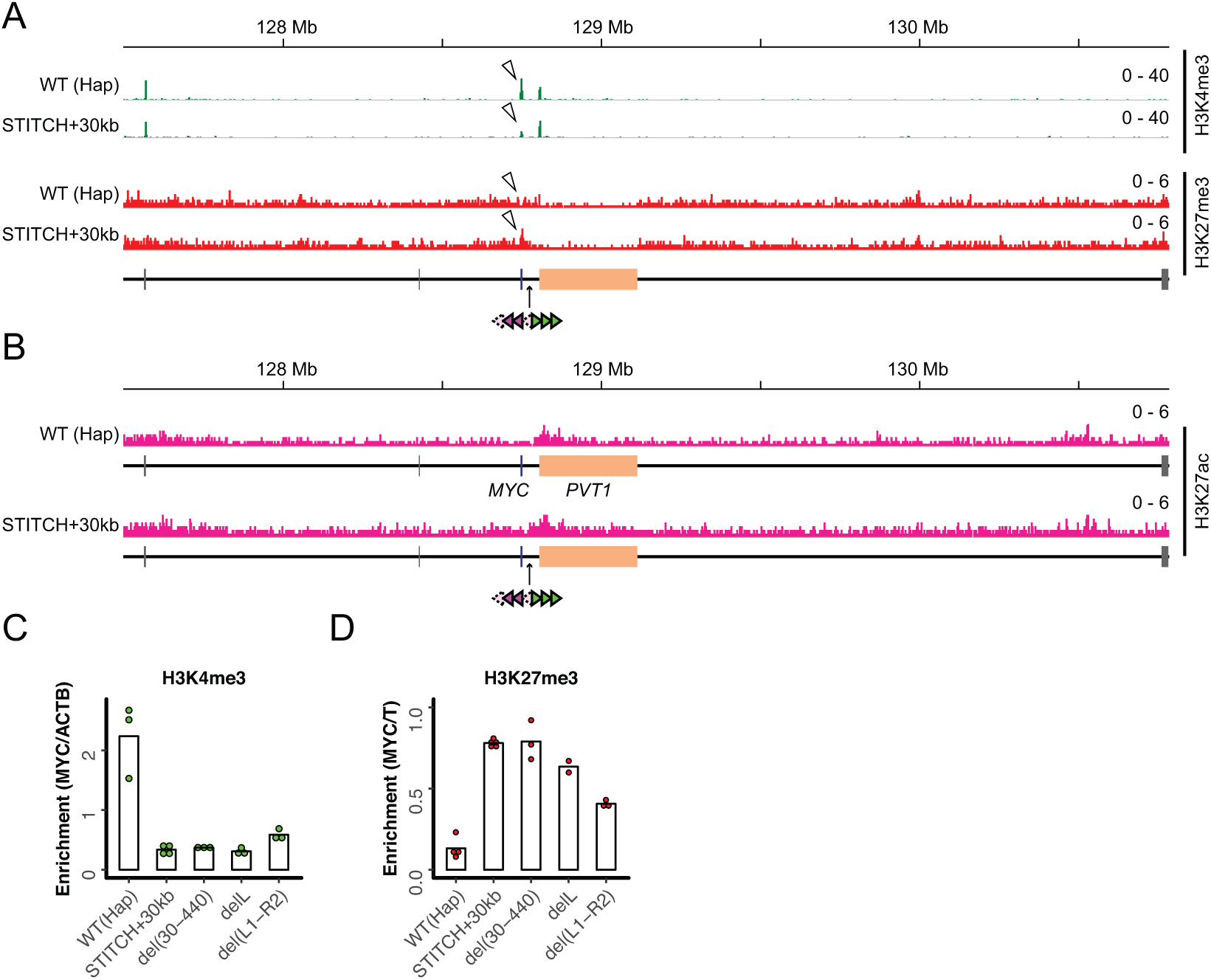
Epigenetic profile around *MYC* with and without STITCH. (A) nChIP-seq for H3K4me3 (green) and H3K27me3 (red) in the wild type (Hap) allele and STITCH+30kb. (B) nChIP-seq for H3K27ac in Hap and STITCH+30kb. (C and D) nChIP-qPCR for H3K4me3 (C) and H3K27me3 (D) in Hap, STITCH+30kb and the indicated mutant alleles of STITCH. The enrichment at *MYC* was normalized with those at *ACTB* (C) and *T* (D). The dots represent replicates, and the bars indicate their means.

**Figure 5.**
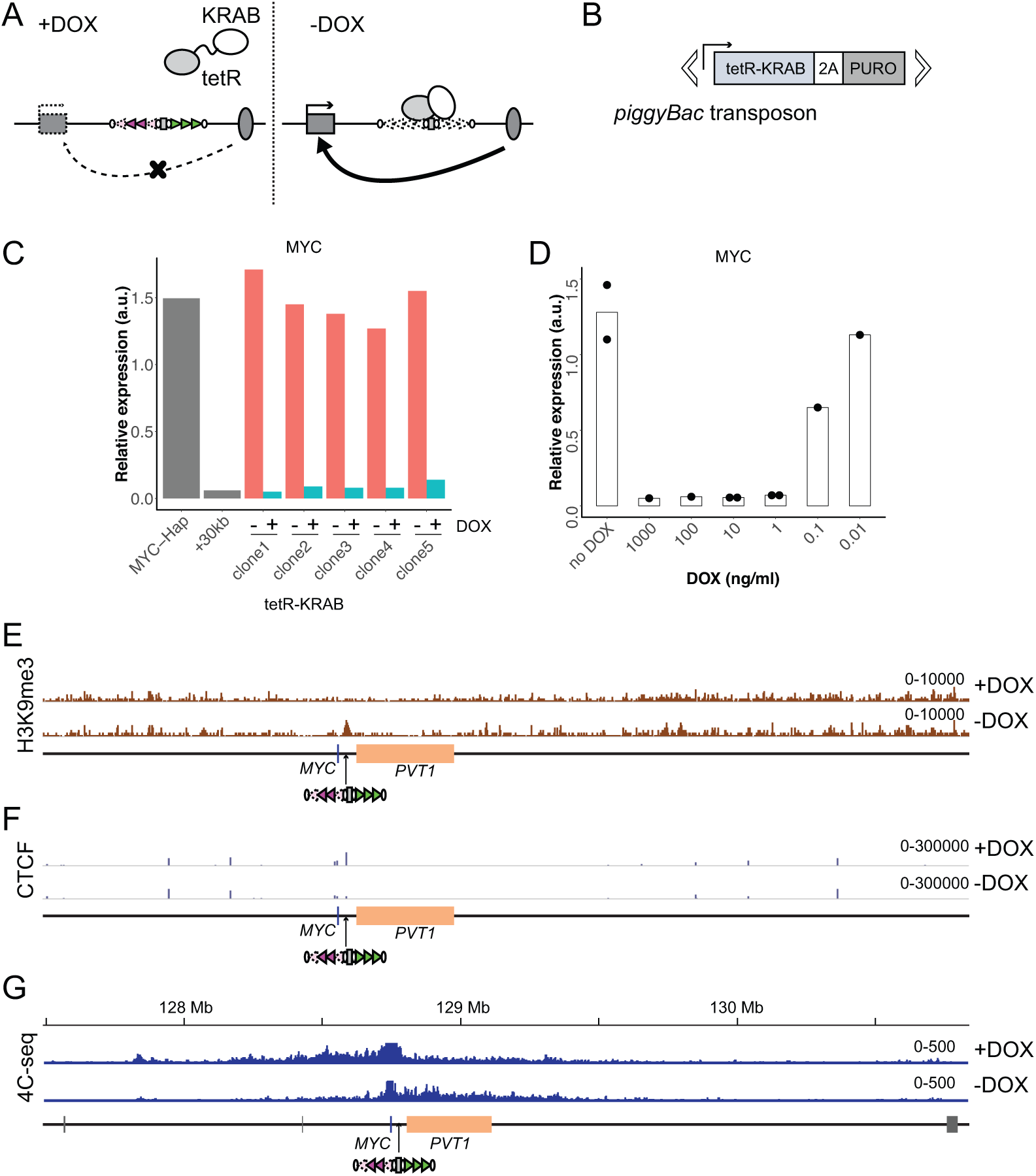
Drug-inducible control of STITCH insulation with tetR-KRAB. (A) DOX dependent binding to and dissociation from STITCH of tetR-KRAB. (B) The piggyBac transposon with the tetR-KRAB transgene followed by a sequence encoding 2A peptide and puromycin resistant gene. (C) The relative expression levels of *MYC* normalized to *ACTB* in five independent clones of STITCH/KRAB with and without DOX were compared to the mean expression levels of Hap and STITCH+30kb clones in Figure 1A. (D) The *MYC* expression level in the clone 1 of STITCH/KRAB with different concentrations of DOX. The dots represent data from replicate experiments and the bars indicate the means. (E, F) nChIP-seq tracks for H3K9me3 (E) and CTCF (F) of the clone 1 with and without DOX. The reads were mapped to a synthetic genomic DNA sequence around the *MYC* locus carrying the STITCH insert. (G) The 4C-seq tracks with and without DOX from VP-MYC2.

### Epigenetic states of *MYC* well correlate with the gene activation by the enhancer

We next investigated how the STITCH insulation of the enhancer impinges on the epigenetic modifications of histones around *MYC*. Active transcription is associated with H3K4me3 at gene promoters, while repressed genes are often marked by H3K27me3. In pluripotent stem cells, many silent developmental genes are marked by the both as so called bivalent states (Bernstein et al., 2006; Mikkelsen et al., 2007; Pan et al., 2007; Zhao et al., 2007). In the wild type allele, *MYC* is exclusively marked by H3K4me3, but not by H3K27me3. Upon the STITCH insulation, the H3K4me3 deposition remained, but was markedly decreased (Figure 4A, Figure S6). Instead H3K27me3 was enriched, and thus became a bivalent state (Figure 4A, Figure S6). By contrast, the neighboring *PVT1* gene has the H3K4me3 at the promoter in the both conditions (Figure 4A, Figure S6). Also the H3K27ac mark around the super-enhancer region was similarly observed (Figure 4B, Figure S6). These results mean that the epigenetic change only occurred at *MYC* upon isolation from the enhancer by STITCH. In the alleles with the STITCH mutations (Figure 3A), the epigenetic states were intermediate between the active and repressive states (Figure 4C and D). Thus, the histone marks around *MYC* vary depending on the association levels with the enhancer and/or the gene expression level.

### Induction of a heterochromatic state by tetR-KRAB impairs the STITCH insulation

The KRAB domain can induce heterochromatin formation around the tetO when linked to tetR (tetR-KRAB) and recruited there (Deuschle et al., 1995; Groner et al., 2010; Sripathy et al., 2006). If this leads to impairment of CTCF bindings as implicated in a previous study (Jiang et al., 2017), it would be possible to control the insulation ability of STITCH by DOX (Figure 5A). To test this, we integrated a transgene consisting of tetR-KRAB followed by DNA encoding the 2A peptide and the puromycin resistant gene with piggyBac transposition into the genome and established several cell lines that stably express it (Figure 5B). The expression levels of the transgene varied much among them (Figure S7A). Nonetheless, in all the cell lines tested, *MYC* expression was repressed in presence of DOX, but became activated after removal of DOX (Figure 5C). When tetR linked to 3xFLAG with HA tag was introduced, *MYC* was not activated without DOX (Figure S7B). Titration of the DOX concentration showed that 1 ng/ml is enough to achieve STITCH insulation in the tested clones with different expression levels of the transgene (Figure 5D and S7C). We performed nChIP-seq for H3K9me3, a mark representing the heterochromatin state, and for CTCF.

When DOX was present, no H3K9me3 peak appeared around the inserted STITCH (Figure 5E, Figure S7D, E); instead, CTCF was strongly bound there (Figure 5F, Figure S7D, E). Accordingly STITCH kept blocking the contacts of *MYC* towards *PVT1* (Figure 5G, Figure S7F). In the absence of DOX, however, H3K9me3 became highly enriched around STITCH (Figure 5E, Figure S7D-F). Concomitantly, the CTCF binding was strongly reduced and the contact of *MYC* well extended to the enhancer region (Figure 5F, G, Figure S7D, E). Thus the STITCH/KRAB system functions as a drug-inducible topological insulator to control gene activation by enhancers.

We next followed temporal changes of the system upon addition and removal of DOX (Figure 6). The nChIP-qPCR for H3K9me3, the 4C-seq assays, and gene expression assays show that 16-24 hours, but not 8 hours, are sufficient to switch the STITCH insulation and *MYC* expression upon both removal and addition of DOX (Figure 6A-E). We tested how the switching of *MYC* expression affects the cell proliferation and found that addition of DOX (i.e. repression of *MYC*) for five days resulted in about 40% reduction of proliferated cells (Figure S7G).

**Figure 6.**
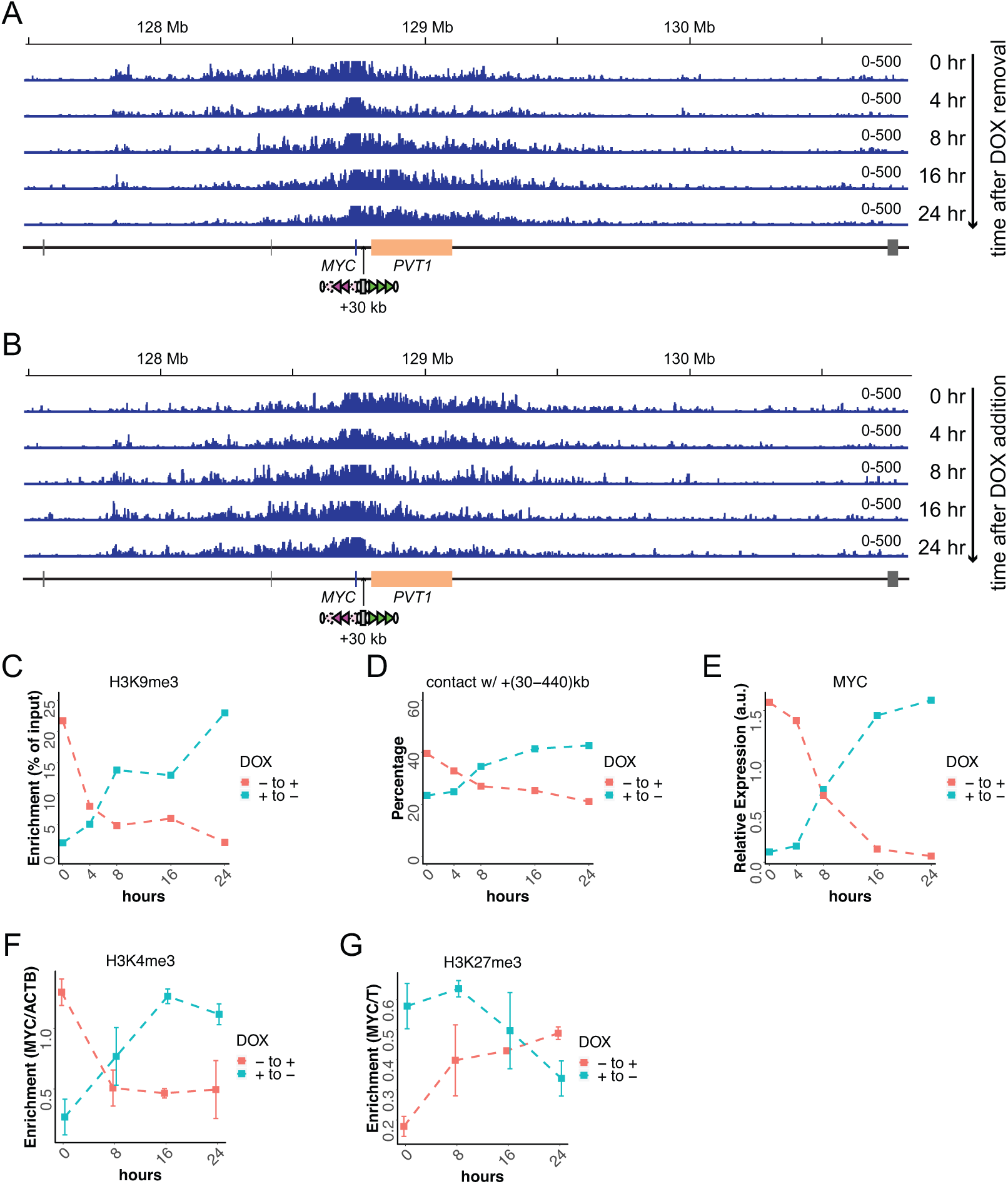
Temporal changes of STITCH insulation upon removal and addition of DOX. (A, B) The 4C-seq profiles in 0, 4, 8, 16 and 24 hours after removal (A) and addition (B) of DOX. (C-G) Temporal changes of nChIP-qPCR for H3K9me3 at STITCH (C), 4C contact frequency with +(30-440)kb region from VP-MYC2 (D), the relative *MYC* expression level normalized to *ACTB* (E), relative enrichment of H3K4me3 at *MYC* normalized with that at *ACTB* (F), and relative enrichment of H3K27me3 at *MYC* normalized with that at *T* (G). The nChIP-qPCR for H3K4me3 and H3K27me3 were performed for three replicate samples. The means and standard deviations (SD) are represented in the plots.

### The epigenetic state of *MYC* follows and reflects the gene expression level

The H3K4me3 and H3K27me3 histone marks correlate well to the gene expression level (Figure 4 and Figure S6). The rapid control of STITCH insulation with KRAB offers us an opportunity to investigate if the epigenetic changes precede the gene expression changes or not. Therefore we also profiled the H3K4me3 and H3K27me3 levels around *MYC* along different time points. Interestingly, while the H3K4me3 mark returned to the levels expected from the gene expression levels within 24 hours after both removal and addition of DOX (Figure 6F), the H3K27me3 did not (Figure 6G). This result suggests that the change of the repressive histone mark follows, but does not precede, the gene expression change.

To test the hypothesis and confirm the reproducibility, we again sampled cells at time points of 24 and 72 hours after addition/removal of DOX. The control samples were cells that were either with or without DOX for more than one passage, respectively. First, we confirmed that the *MYC* expression was up- and down-regulated within one day after removal and addition of DOX to the levels of the controls, respectively (Figure 7A and B). Then we performed nChIP-qPCR for the both histone marks. Consistently to above, the deposition of H3K27me3 was significantly higher and lower in 24 hours than 72 hours and controls after removal and addition of DOX, respectively (Figure 7C and D). By contrast, we did not see such significant differences for H3K4me3, suggesting that the active mark is more rapidly turned over than the repressive mark (Figure 7E and F).

**Figure 7.**
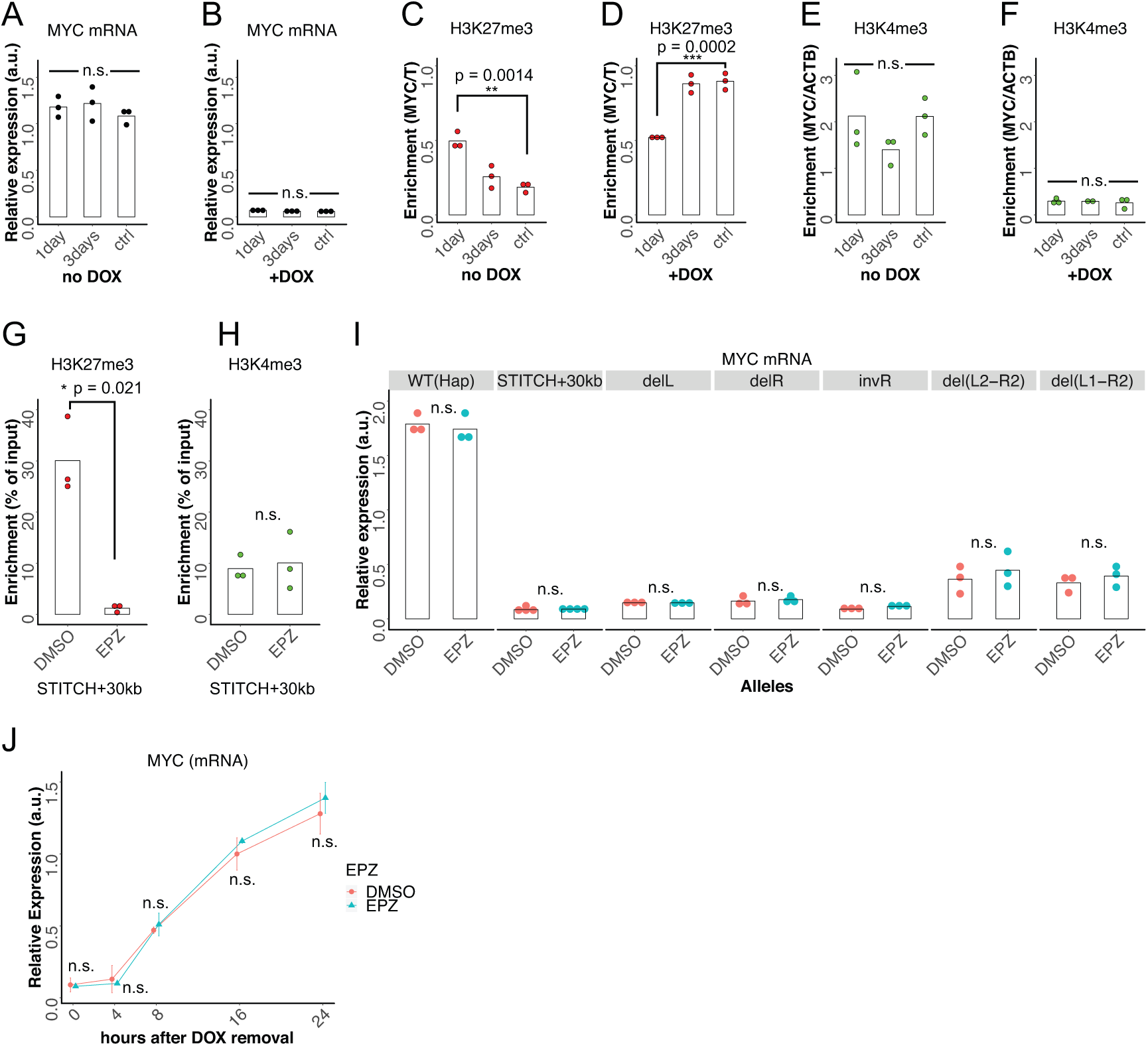
Delayed turnover of H3K27me3 enrichment after the gene expression change. (A-F) Relative *MYC* expression level normalized to *ACTB* (A and B), relative H3K27me3 enrichment at *MYC* normalized to *T* (C and D) and relative H3K4me3 level at *MY*C normalized to *ACTB* (E and F) were measured after 24 hours (1day), 72 hours (3days) and longer than one passage (ctrl) from either removal (A, C, E) or addition (B, D, F) of DOX in the STITCH/KRAB. The dots represent data of replicates, and the bars indicate their means. **, *** and n.s. indicate p < 0.01, p < 0.001 and p > 0.05, respectively, by one-way ANOVA. The p-values with Tukey’s multiple-comparison post hoc test are indicated. (G, H) Enrichment of H3K27me3 (G) and H3K4me3 (H) at *MYC* after two days treatment with EPZ or DMSO. The dots represent replicates, and the bars indicate their means. (I) Relative *MYC* expression levels in the Hap, STITCH+30kb and the mutants of STITCH after three days treatment of EPZ or DMSO. The dots represent replicates, and the bars indicate their means. (J) Temporal changes of relative *MYC* expression levels after DOX removal in the STITCH/KRAB. Before DOX was removed, cells were exposed to EPZ or DMSO for two days. Means and SD of three replicate experiments were plotted. * and n.s. indicate p < 0.05 and > 0.05, respectively, by Welch’s two sample t-test in (G-J).

These results suggest that the H3K27me3 mark par se only reflects, but does not determine the gene expression level. To test this, we treated the cells with EPZ-6438 (EPZ), an inhibitor of Enhancer of zeste homolog 2 (EZH2), an enzymatic subunit of Polycomb Repressive Complex 2 (PRC2), which catalyzes methylation of H3K27 (Knutson et al., 2013). Addition of the inhibitor at 200 nM for 2 days was enough to mostly diminish the H3K27me3 mark around the *MYC* genic region (Figure 7G). This reduction of H3K27me3 did not result in significantly higher enrichment of the active H3K4me3 mark (Figure 7H). We compared the *MYC* expression levels in Hap, STITCH+30kb, and the mutant alleles of STITCH treated with EPZ or DMSO for 3 days (Figure 7I). The difference between the two treatments was not significant in any of the alleles. We next treated the STITCH/KRAB cells with EPZ or DMSO for two days, removed DOX, and compared the *MYC* expression at different time points up to 24 hours after removal of DOX. The expression profiles showed no significant difference between the two, suggesting that the H3K27me3 mark does not affect the gene activation by the enhancer (Figure 7J).

## DISCUSSION

### STITCH as a novel tool for manipulating gene expression

STITCH blocks interaction of genes and enhancers when inserted in between as an insulator element. Further combining this with the DOX control of tetR-KRAB achieved drug-inducible switching of the insulation. Thus the system adds a new layer to the toolkits for manipulating gene expression.

In this study, we show its utility in studying gene regulation by long-range enhancers, which in fact led to several important findings as described above and discussed below. The uniqueness of the system in this respect is that it can target specifically only one locus without affecting much of the cellular states and the epigenome even around the enhancer region (Figure 4 and Figure S6). The changes of gene expression and epigenetic states observed at *MYC* should be mostly attributed to the association or dissociation with the enhancer controlled by the STITCH insulation, but not to effects in the genome-wide or cellular level. This is in contrast to many other studies that depleted genes and proteins or induced cellular differentiation and signaling cascades. Coupling the STITCH/KRAB system with live-imaging technique and so on should further contribute to understanding gene regulation by enhancers.

We also anticipate that STITCH can be an important tool to disrupt gene function in a tissue-specific manner. Currently this is predominantly achieved by the Cre-loxP system, which inevitably needs a good driver for Cre expression. However, STITCH disruption needs just one insertion between a gene and an enhancer. We expect that this should work even if the enhancer stretches over a wide region like a super-enhancer as exemplified in this study. Controlling the insulation by KRAB can add another degree of control.

### Mechanism of the STITCH insulation and its control by heterochromatin induction

Though CTCF binding to L1 and L4 was not confirmed by the nChIP, the other five sites were directly bound by CTCF (Figure S1C and Figure S7E). The delL and del(L1-R2) alleles only keep the direct binding sites of CTCF, and still show substantial insulation activity (Figure 3). Further, the insulation activity and the folding property is dependent on the orientation of the binding motifs (Figure 3), as in the endogenous TZ region (Tsujimura et al., 2018). Therefore, it should be safe to attribute the STITCH insulation primarily to the bindings of CTCF. Blocking of enhancer activity by heterologously inserted CTCF binding sites is consistent with previous studies (Bell et al., 1999; Liu et al., 2015).

It is not clear, however, why CTCF disrupts gene activation by enhancers so much. CTCF generally impairs contacts between the flanking regions together with cohesin, and often establishes a boundary of contact domains (Nora et al., 2017). Numerous studies have shown that the domain boundaries insulate enhancer activation (Dowen et al., 2014; Lupiáñez et al., 2015; Narendra et al., 2015; Symmons et al., 2014; Tsujimura et al., 2015; 2018). From these observations, it is vaguely accepted that CTCF restricts chromatin contacts, which leads to limitation of enhancer association within contact domains. However, contact frequency is not so drastically reduced as gene expression levels by presence of CTCF bindings. For example, STITCH at most reduces the contacts of *MYC* with the +(30-440)kb region by half, but the decrease of gene expression exceeds 20 folds (Figure 1, Figure S1). Also, a recent comprehensive imaging of chromatin structure at given loci showed that the domain-like structures are frequently present beyond the boundary positions (Bintu et al., 2018). Therefore, it is not likely that the disruption of gene activation is only attributed to the simple reduction of the contacts beyond the domain boundary. The escalated association of *MYC* with the super-enhancer/*PVT1* region upon the stepwise loss of CTCF binding sites of STITCH might be suggestive in this respect (Figure 3). This observation well explains, at least in part, the gradual change of contact frequency is translated to the skewed expression changes. The fact that loss of CTCF or cohesin induces association among active regions (Nora et al., 2017; Rao et al., 2017; Schwarzer et al., 2017) suggests that the preferential association of *MYC* with the super-enhancer also obeys the same compartmentalization principle. Then how does the CTCF binding interrupt this process? Possibly there might be a mechanism that boosts aggregation of the active regions upon increase of contact frequencies. For example, the recently proposed phase separation model may explain it well (Hnisz et al., 2017). In addition, or alternatively, CTCF *per se*, perhaps through loop extrusion and/or stabilization by cohesin, disrupts the aggregation, when present in between. Uncoupling these different effects would be challenging.

The induction of tetR-KRAB impaired binding of CTCF at STITCH and restored the contacts of *MYC* with the enhancer over STITCH. We think this is due to the formation of the heterochromatic states that were represented by the H3K9me3 deposition. The heterochromatic regions form dense nucleosomes, which may exclude binding of transcription factors (Machida et al., 2018). A previous study already suggested that KRAB induction reduces binding of CTCF (Jiang et al., 2017). It is notable that the formation of heterochromatin does not prevent association between genes and enhancers. At the same *MYC* locus, in fact, recruitment of the KRAB domain to the *PVT1* promoter did not block the enhancer activation (Cho et al., 2018).

### The *MYC* regulation

*MYC* is one of the four factors of the original cocktail to induce pluripotent stem cells (Takahashi and Yamanaka, 2006; Takahashi et al., 2007). In the STITCH/KRAB cells, the decrease of *MYC* expression led to decreased proliferation rate (Figure S7F). Further our transcriptome analysis revealed that down-regulation of *MYC* leads to decrease of genes involved in cholesterol synthesis and ribosomal assembly (Figure 2). Genes related to ribosomes have been well known targets of *MYC* in various cell types, and are thought to contribute to increasing cell proliferation (Hofmann et al., 2015; Pulikkan et al., 2018; Uslu et al., 2014; van Riggelen et al., 2010; Zeller et al., 2006). Also *MYC* haploinsufficient mice showed reduction of cholesterol synthesis through down-regulation of genes involved in the process (Hofmann et al., 2015). Cholesterol is highly relevant to promoting cell proliferation in stem cells and cancers as a major component of cell membranes, and possibly as a mitogen (Wang et al., 2018). Further, cholesterol is important for mitochondrial function (Ribas et al., 2016), and thus seems to have a role in energy metabolism of stem cells and cancer, where glycolysis is more dominant than mitochondrial respiration (Gu et al., 2016). These facts suggest that in the iPSCs *MYC* regulates cellular metabolism and proliferation through up-regulation of a certain set of genes that are also shared by different types of cells including cancer cells. Further digging into the function of *MYC* in our system should be fruitful in this sense.

*MYC* expression is regulated by tissue-specific enhancers, in contrast to the generality of its function. Tight regulation by enhancers should be critical for organismal development and homeostasis considering the roles of *MYC* in the cellular processes. So far, many enhancers have been identified around the locus in different cell types including cancer cells (Bahr et al., 2018; Cho et al., 2018; Dave et al., 2017; Herranz et al., 2014; Hnisz et al., 2013; Lovén et al., 2013; Pulikkan et al., 2018; Shi et al., 2013; Sur et al., 2012; Uslu et al., 2014; Zhang et al., 2016). Their locations vary much. Some are distant and others are similarly embedded in the *PVT1* genic region as in the iPSCs. It should be noted that *PVT1* is annotated as a lncRNA gene. Although it has been proposed that *PVT1* has functions as lncRNA particularly in cancer cells, the molecular mechanism remains elusive (Cui et al., 2016). A recent study shows that inhibition of the *PVT1* transcription does not impact on *MYC* expression in a cancer cell line (Cho et al., 2018). The study rather showed that the *PVT1* promoter modulates *MYC* expression as a competitor for enhancer activity (Cho et al., 2018), which may indicate the transcribed RNA is a byproduct. In our study, del(30-440) exhibited a rather milder effect on the transcriptome than STITCH+30kb (Figure 2E and F). These data may favor a notion that *PVT1* has little function as lncRNA, if any.

### The H3K27me3 mark reflects the gene expression

The STITCH insulation not only down-regulated the gene expression but also affected the epigenetic states of *MYC*. At the normal state where *MYC* is highly expressed, the promoter is marked by H3K4me3, but not by H3K27me3. When the enhancer activity was blocked, the gene expression was repressed and the prominent H3K27me3 histone mark appeared (Figure 4). The deletions and inversion of the CTCF binding sites in STITCH led to the intermediate states (Figure 4C and D). These results corroborate the well-established correlation of gene expression levels and the epigenetic states (Hosogane et al., 2016; Højfeldt et al., 2018; Narendra et al., 2015; Riising et al., 2014).

We further investigated the temporal change and showed that deposition of H3K27me3 only follows and reflects, but does not precede and affect gene expression changes. Perhaps this might seem contradictory to the prevailing notion of the histone mark as a repressor. However, the delayed change of the histone modification after transcriptional change is consistent with previous reports showing the same relationship upon global induction of cellular stimuli (Hosogane et al., 2013; Kashyap et al., 2011), and with the mechanistic property of the repressive state as an epigenetic memory (Reinberg and Vales, 2018). Also accumulative evidence has shown that PRC2 has almost no effect on gene expression in a certain context (Riising et al., 2014). Yet, mutations in genes encoding PRC2 components have indicated PRC2 has diverse and critical roles in organisms (Schuettengruber and Cavalli, 2009). Also it was shown that PRC2 maintains gene silencing during differentiation of mouse ES cells (Riising et al., 2014), To explain these observations, it has been proposed that deposition of H3K27me3 raises the threshold for gene activation (Comet et al., 2016). However, the studies involving gene activation so far were carried out under global induction of cellular stimuli. Therefore, it has not been clear if the H3K27me3 marks regulate gene activation locally as a resistance in *cis* or rather globally through effects on the cellular and epigenomic states. In fact, our experiment showed presence of H3K27me3 makes no significant difference in *MYC* activation upon the local induction by the enhancer (Figure 7J), and thus challenged the above hypothesis. The role of this repressive histone mark needs to be further studied in future.

## Acknowledgements

We thank Prof. Shinya Yamanaka for providing us the hiPSC line. This work was supported by research grants from Mutou Group (http://www.wism-mutoh.co.jp); APA Group (https://www.apa.co.jp); IMS Group (http://www.ims.gr.jp/group/); Alba Lab (http://www.albalab.co.jp), and Kobe One Medicine, One Health (http://kobe.omoh.jp), as well as by Grants-in-Aid for Young Scientists (B) (No. 17K16072 to TT), for Scientific Research (B) (No. 15H03001 to KH) and for Scientific Research (C) (Nos. 16K09602 to OT, and 15K09244 to MY) from the Japan Society for the Promotion of Science. The funders had no role in the design of the study and in writing the manuscript.

## Authors’ contributions

Taro Tsujimura and Keiichi Hishikawa conceived the study. OT, YM, Kazuto Hoshi, Tsuyoshi Takato and Keiichi Hishikawa helped establishing a system to culture the iPS cells and provided resources for it. Taro Tsujimura and ES performed the cell culture, genome editing, RNA extraction, processing of 4C samples, nChIP and qPCR assays. Taro Tsujimura prepared the libraries for deep sequencing. AT performed the deep sequencing and curated the data. Taro Tsujimura analyzed the deep sequencing data. MH, Kazuto Hoshi, Tsuyoshi Takato, HO and Keiichi Hishikawa administered the experiments and supported the project. Taro Tsujimura, HO and KH wrote the manuscript with inputs from the other authors.

## Availability of data and materials

All the deep sequencing data of the 4C-seq, RNA-seq and nChIP-seq libraries analyzed in this study were deposited in ArrayExpress.

## Materials and methods

### Cell culture

The human iPSC line 253G1 (Nakagawa et al., 2007) was kindly provided by Prof. Shinya Yamanaka. We cultured the cells in the StemFit^®^ AK02N medium (ReproCELL, Cat#RCAK02N) on dish coated with iMatrix-511 (ReproCELL, Cat#NP892-012) without feeder cells. We added Y-27632 (FUJIFILM Wako, Cat#036-24023) at the final concentration of 10 μM, when seeding the cells on dish. We used the 0.5x of TrypLE™ Select (Thermo Fisher Scientific K.K., Cat#12563-011) to dissociate the cells for passaging.

DOX (Sigma, Cat#D9891) was basically added at the final concentration of 10 ng/ml, unless specifically indicated. When DOX was removed for time-course analysis, the concentration was first changed to 1 ng/ml one day before the start of removal. Then at the start of the removal, the cells were first washed with PBS (Thermo Fisher Scientific K.K., 10010-049) and then fresh medium without DOX was supplied. Further two hours later, wash with PBS and replacement of medium was repeated to ensure the removal of DOX. EPZ (Adipogen Life Sciences, Cat#SYN-3045-M001) was used at the final concentration of 200 nM. For the DMSO controls, same volume of DMSO as EPZ was added.

### Genome editing

To delete the 3-Mb region of the *MYC* locus, we co-transfected RNP complex of CRISPR/Cas9 targeting the both edges of the deletion interval with Lipofectamine™ RNAiMAX Transfection Reagent (Thermo Fisher Scientific K.K., Cat#13778030). We assembled the RNP from Alt-R™ CRISPR crRNA (Integrated DNA Technologies, listed in Table S1), Alt-R™ CRISPR tracrRNA, and Alt-R™ S.p. Cas9 Nuclease 3NLS (Integrated DNA Technologies, Cat#1072532 and Cat#1074181, respectively), following the manufacturer’s protocol. The target sequences of the guide RNAs are described in Table S1. After the transfection, cells were sparsely re-plated on dish. Grown colonies were picked up and expanded. The clones were screened for the correctly edited allele by PCR genotyping (see Table S4 for the primer sequences). We then confirmed the deletion by direct Sanger sequencing.

The STITCH vector targeting into the +30kb position with the homology arm of 150-bp length at each side was synthesized by Integrated DNA Technologies (see Table S2 for the DNA sequences). We amplified the fragment by PCR (see Table S3 for the primer sequences) with Tks Gflex™ DNA Polymerase (Takara, Cat#R060A) and purified it. Then we transfected it into the cells with Lipofectamine™ 3000 Transfection Reagent (Thermo Fisher Scientific K.K., Cat#L3000001) together with a RNP complex of CRISPR/Cas9 targeting the insertion site as described above and transfected with Lipofectamine™ RNAiMAX Transfection Reagent (Thermo Fisher Scientific K.K., Cat#13778030). See Table S1 for the target sequences of the guide RNAs. The positive cells were first selected in the culture medium with 0.2 mg/L puromycin. Then survived colonies were picked up and expanded. The correct insertion was confirmed by PCR and direct sequencing. We found a single nucleotide mutation within the R3 sequence in the clone that we obtained, which was far away from the core motif for CTCF binding for more than 30 bp. To insert STITCH into the other four sites, we attached 50-bp homology arms by PCR using the STITCH vector as the template, and performed the transfection as the same way as above. We screened puromycin resistant clones and then confirmed the insertion by PCR. These targeted cells were further transfected with Cre Recombinase encoding mRNA (OZ Biosciences, Cat#MRNA32-20) using Lipofectamine™ RNAiMAX Transfection Reagent (Thermo Fisher Scientific K.K., Cat#13778030) to remove the puromycin resistant cassette (Figure 1A). After the transfection, the cells were sparsely plated on dish, and colonies were picked up after they formed. We screened positive clones by PCR (see Table S4 for the primer sequences).

To delete or invert the CTCF binding sites within STITCH, we transfected CRISPR/Cas9 RNPs targeting the edges of the intervals of the deletion/inversion as described above (see Figure S2B-E). The target sequences of the guide RNAs are described in Table S1. After transfection with the RNPs, the cells were sparsely seeded and grown colonies were picked up. The mutations were first screened by PCR (see Table S4 for the primer sequences). Then the DNA sequences were confirmed by direct sequencing. While we tried to obtain the del(L2-R2) clones, we obtained the del(L1-R3) clone probably due to the excessive excision at the cutting site (Figure S2D).

To make the del(30-440) allele, we targeted the selection cassette only (i.e. the two loxP sites sandwiching the Puromycin resistant gene inside) of the STITCH vector into the +440kb position of a delL clone in the same way as above (Figure S2A). The targeting fragment was prepared by two rounds of PCR from the STITCH vector (see Table S3 for the primer sequences). After correct integration, Cre Recombinase encoding mRNA (OZ Biosciences, Cat#MRNA32-20) was transfected, and the deletion allele was selected by PCR screening (see Table S4 for the primer sequences).

To obtain cells that stably express tetR-KRAB, we designed a plasmid vector of a piggyBac transposon carrying coding sequence for tetR-KRAB followed by that of the 2A peptide and the puromycin resistant gene under the promoter of human PGK gene. The plasmid was synthesized by GenScript. We also designed the piggyBac vector containing tetR-3xFlag-HA and obtained the plasmid synthesized by GenScript. We transfected the plasmids with Super PiggyBac Transposase Expression Vector (System Biosciences, PB210PA-1) using Lipofectamine™ 3000 Transfection Reagent, and screened positive clones under puromycin selection, as described above. The positive cells were expanded and maintained in presence of puromycin at 0.1 mg/L. We obtained and characterized several clones, but picked one (the clone 1 in Figure 5A) for the subsequent analysis of STITCH/KRAB.

### RNA extraction, cDNA synthesis, qPCR and library preparation for RNA-seq

RNA was extracted using the High-pure RNA isolation kit (Roche, Cat#11828665001) in presence of the DNase I included in the kit. We subsequently synthesized the cDNA with the High-Capacity cDNA Reverse Transcription Kit (Thermo Fisher Scientific K.K., Cat#4368813). We used KAPA SYBR Fast qPCR Kit (Kapa Biosystems, Cat#KK4621) as the reagent and the Applied Biosystems^®^ 7500 Fast Real-Time PCR System (Thermo Fisher Scientific K.K.) for the qPCR reaction. The primers used for qPCR assays are listed in Table S5. To prepare libraries for RNA-seq, we first enriched mRNA using NEBNext^®^ Poly(A) mRNA Magnetic Isolation (New England Biolabs, Cat#E7490S). Then subsequently, we used NEXTflex Rapid RNA-Seq kit (Bioo Scientific, Cat#NOVA-5238-01) for the library preparation with customly designed oligo DNAs (listed in Table S6) as primers for the PCR reaction. The libraries were sequenced with HiSeq2500 System (Illumina) using HiSeq SR Rapid Cluster Kit v2-HS (Illumina, Cat#GD-402-4002) and HiSeq Rapid SBS Kit v2-HS 50 Cycle (Illumina, Cat#FC-402-4022).

### 4C-seq library preparation and sequencing

For a 4C-seq library prep, we collected c.a. 1 million cells and fixed them in 2% paraformaldehyde for 10 minutes at room temperature. Then the cells were lysed in lysis buffer (50 mM Tris (pH7.5), 150 mM NaCl, 5 mM EDTA, 0.5% NP-40, 1% Triton X-100, 1x complete proteinase inhibitors (Roche, Cat#11697498001); 1 ml), passed through a 23-guage needle, pelleted and frozen in liquid nitrogen. After the cells were resuspended in H_2_O and CutSmart^®^ Buffer (New England Biolabs, Cat#B7204) and treated with 0.3% SDS and 2.5% Triton X100 at 37°C for 1 hour, respectively, we performed first digestion of the chromatin with 25 units of *Nla*III restriction enzyme (New England Biolabs, Cat#R0125) on a rotator at 37°C for overnight. After heat inactivation of the enzyme, 12.5 units of T4 DNA ligase (Thermo Fisher Scientific, Cat#EL0014) were applied for self-ligation of the digested chromatin. After de-crosslinking and purification, we carried out second digestion with 20 units of *Dpn*II restriction enzyme (New England Biolabs, Cat#R0543). Then the chromatin was again self-ligated with 12.5 units of T4 DNA ligase (Thermo Fisher Scientific, Cat#EL0014). We then performed the inverse PCR from the chromatin of the c.a. 1 million cells as the template to amplify the 4C library from a given viewpoint for 25 cycles with Tks Gflex™ DNA Polymerase (Takara, Cat#R060A). The primer sequences for VP-MYC1 were 5’-TCTTTCCCTACACGACGCTCTTCCGATCTGTAGGCGCGCGTAGTTAATTC-3’ and 5’-GTGACTGGAGTTCAGACGTGTGCTCTTCCGATCTTCGCTAAGGCTGGGGAAAG-3’; those for VP-MYC2 were 5’-TCTTTCCCTACACGACGCTCTTCCGATCTTCTCCCTGGGACTCTTGAT-3’ and 5’-GTGACTGGAGTTCAGACGTGTGCTCTTCCGATCTAAACTCCCATTGCATTTGTTG-3’. We purified the DNA with High-pure PCR Product Purification Kit (Roche, Cat#11732676001) and performed the 2nd round of PCR to attach to the libraries adaptor and index sequences for the NGS analysis for 8 cycles again with Tks Gflex™ DNA Polymerase (Takara, Cat#R060A). The DNA sequences of the adaptor/index primers are listed in Table S6. The DNA was purified with High-pure PCR Product Purification Kit (Roche, Cat#11732676001). The final libraries were pooled and sequenced with the HiSeq2500 system as described above. Note that the sequences were read from the *NlaIII* side for VP-MYC1 and the *DpnII* side for VP-MYC2.

### nChIP for histone modifications and CTCF binding, qPCR, and library preparation for sequencing

To perform nChIP for histone modifications, cells were dissociated from dish with TrypLE Select (Thermo Fisher Scientific K.K., Cat#12563-011), washed with PBS, and frozen as pellets. After resuspension in ChIP dilution buffer (20 mM Tris-HCl pH8.0, 150 mM NaCl, 2 mM EDTA, 1% Triton X-100), supplemented with 0.05% SDS, 3 mM CaCl_2_ and protease inhibitors, they were incubated on ice for 10 minutes, and incubated at 37°C for 2 minutes. We added 0.48 μl of micrococcal nuclease (NEB, Cat#M0247S) per 1.0 million cells, and incubated them at 37°C for 10 minutes. To stop the digestion reaction, EDTA and EGTA were added so the final concentration was 10 mM and 20 mM, respectively. To solubilize the chromatin, we applied sonication with Ultrasonic Homogenizer UH-50 (SMT Co., Ltd.) for three times of 20-seconds pulse and incubated them at 4°C for 1 hour. The solubilized chromatin after removal of the cell debris by centrifugation was incubated with antibody at 4°C for overnight. We used 0.6, 0.4, 0.5 and 0.6 μl of antibodies per 400,000 cells for H3K4me3, H3K27me3, H3K9me3 and H3K27ac (MAB Institute, Cat#MABI0304S, Cat#MABI0323S, Cat#MABI0318S and Cat#MABI0309S), respectively. The chromatin with the antibodies was incubated with 6 μl of Dynabeads Protein G (Thermo Fisher Scientific, Cat# 10003D) for one hour. Then the beads were washed for three times with ChIP dilution buffer supplemented with 0.05% SDS, and subsequently twice with high-salt wash buffer (20 mM Tris-HCl pH8.0, 500 mM NaCl, 2 mM EDTA, 1% Triton X-100, 0.05% SDS). The chromatin was treated with RNaseA (50 ng/μl) at 37°C for 15 minutes and then with Proteinase K (100 ng/μl) at 55°C for 1 hour in ChIP extraction buffer (20 mM Tris-HCl pH 8.0, 300 mM NaCl, 10 mM EDTA, 5mM EGTA, 0.1% SDS). The DNA was precipitated with ethanol and eluted in 10 mM Tris-HCl pH 8.0 after removal of the beads. We performed the CTCF nChIP exactly as described before with the same polyclonal anti-CTCF antibody (Millipore, Cat#07-729) (Tsujimura et al., 2018). For qPCR assays, we used KAPA SYBR Fast qPCR Kit (Kapa Biosystems, Cat#KK4621) as the reagent and the Applied Biosystems^®^ 7500 Fast Real-Time PCR System (Thermo Fisher Scientific K.K.) as the platform. To prepare nChIP-seq libraries, we used the NEBNext Ultra II DNA Library Prep with Sample Purification Beads (NEB, Cat#E7103S). We basically followed the protocol from the manufacturer, but used partly oligo DNAs that were customly designed by us for the PCR reaction as listed in Table S6 and S7. The libraries were sequenced with the HiSeq2500 as described above.

### Data analysis of qPCR assay for gene expression levels

We first measured the amplification efficiency of primer pairs for *MYC*, *ACTB* and the puromycin resistant gene, and confirmed it is nearly 100% for the all. Therefore, we used ΔΔCt method to obtain the relative expression levels normalized to *ACTB*. As a reference sample, we used a large stock of cDNA prepared from the same iPSC line (253G1), which were cultured in a different condition from the present study (with feeder cells in a different medium), and always placed the reference sample in duplicates or triplicates in the same PCR plates, when measuring the Ct values of samples.

Replicates were defined differently for different experimental purposes. For STITCH insertions and del(30-440), replicates mean independent clones that were segregated after Cre transfection. The relative expression levels were measured for each clone and plotted in Figure 1D. For mutant clones of STITCH, replicates mean independent clones after CRISPR/Cas9 genome editing. The relative expression levels were measured for each clone and plotted in Figure 3A. We also obtained sub-clones from Hap and treated them as replicates in Figure 1D and Figure 3A. In the Figures 1A and 3D, mean values of the replicates were also represented as bars. The relative expression levels of the STITCH mutants and Hap clone in Figure 3I and J, and Figure S4B and C were the mean values of the replicates. We obtained five and three clones after transfection of tetR-KRAB and tetR-3xFlag-HA transposons, respectively. The relative expression levels of *MYC* and the puromycin resistant gene were assayed for all of these clones in Figure 5C and Figure S7A and B. For the treatment with DOX and EPZ, we used only one representative clone (the clone 1) for each, and performed replicate experiments, which mean samples separately treated with drugs in different dish (Figures 5D, 7A, B, I and J). We performed one-way ANOVA with Tukey’s multiple-comparison post hoc test to infer statistical significance between different conditions in Figure 7A and B, and Welch’s two sample t-test in Figure 7I and J for the statistical significance between the DMSO and EPZ treatments. The data were represented as graphs with the ggplot2 package in R.

### Cell proliferation assay

To compare cell proliferation rates between conditions with and without DOX, equal volumes of cells were first seeded in three replicates for each from a single population in the same medium without DOX. On the next day, we replaced the medium with fresh one with or without DOX. After five days, the cell numbers were counted using a hemocytometer. We represented the relative proliferation rates as normalized cell numbers divided by the mean number of cells in the DOX minus condition. The assay was performed for both the Hap and STITCH/KRAB clones. We performed Welch’s two sample t-test to infer the statistical significance between the two conditions.

### Data analysis of RNA-seq

We prepared and sequenced libraries from three replicate clones (see above) for each of Hap, STITCH+30kb and del(30-440). We first combined separately sequenced reads of same libraries from different lanes as fastq files. We mapped the sequences to the human genome (hg19) with HISAT2 (Kim et al., 2015). We made BedGraph tracks with HOMER (Heinz et al., 2010) and visualized them in Integrative Genomics Viewer (version 2.4.6) (IGV) (Robinson et al., 2011). The data ranges are indicated by counts per 10 million. We assigned the mapped reads to annotated genes with HTSeq (Anders et al., 2015). We normalized counts and called differentially expressed genes between different conditions with DESeq2, with a threshold of the adjusted p value < 0.1 (Love et al., 2014). We applied the shrunken log2 fold changes for visualization as the MA-plots. The Venn diagram was drawn with the VennDiagram package in R (Chen and Boutros, 2011). The GO term enrichment analysis was performed with the topGO package in R (Alexa et al., 2006). Fisher’s exact test was employed for the statistical test. The data were visualized with the ggplot2 package in R.

### Data analysis of 4C-seq

We only employed a representative clone for each genomic configuration for the 4C-seq assays. However, we prepared a couple of replicate libraries from same clones for each viewpoint, which were separately prepared from different dish, to confirm the reproducibility of the experiment (Figure S1 and S4).

We first combined separately sequenced reads of same libraries from different lanes as fastq files. The sequences of the viewpoint fragment up to the restriction sites were removed with FASTX-Toolkit. Then we mapped the rest of the sequences to the human genome (hg19) using Bowtie2 mostly with the default settings except that the –score-min option was set as “L,-0.1,-0.1” (Langmead and Salzberg, 2012). The generated SAM files were converted to BAM files, indexed and sorted with SAMtools (Li et al., 2009). We used the FourCSeq package to normalize the counts as reads per million (RPM), smooth them with the window size of seven fragments, and produce BedGraph files (Klein et al., 2015). We visualized the tracks in Integrative Genomics Viewer (version 2.4.6) (Robinson et al., 2011). The data ranges are indicated by counts per million. Counting number of reads mapped to given regions was performed with BEDTools (version 2.26.0) (Quinlan and Hall, 2010). To calculate contact frequencies, we divided the read numbers in a given region by the total read numbers mapped to the locally haploid 3-Mb region except the 10-kb region from the viewpoint fragment. To perform PCA, we first counted reads in the 30-kb bins. We combined the read numbers of replicates from the same viewpoint (either VP-MYC1 or VP-MYC2). Then we calculated ratios of reads in each bin within the region of interest (whole locus, the left 900-kb region, or the right 600-kb region). Then we performed PCA using the data sets of VP-MYC1 and VP-MYC2 for each clone with the prcomp function in R. The component loadings were calculated using the sweep function in R. To perform the correlative analysis between the 4C-seq counts and gene expression levels, we also combined reads of replicates and calculated contact frequencies first. Then, the linear regression was performed against the log-log plot to obtain the slope in R. The Spearman’s rank correlation coefficients were also calculated using a function in R. The log-log plots were visualized using the ggplot2 package in R.

### Data analysis of nChIP-qPCR assay

We always took input samples for every nChIP, and calculated enrichment as ratios to the input samples. Our replicates mean different nChIP samples derived from separately cultured cells in different dish. In order to cancel the inevitable variance in the total enrichment efficiency of nChIP experiments, we basically normalized the enrichment at *MYC* to those at control regions, which were the *ACTB* region for the active H3K4me3 mark and the *T* region for the repressive H3K27me3 mark. As the treatment with EPZ causes an epigenetic change in genome-wide, we did not do the normalization in Figure 7G and H. To test statistical significance between different conditions in Figure 7C-F, we performed one-way ANOVA with Tukey’s multiple-comparison post hoc test. To assess statistical significance between treatments with DMSO and EPZ, we performed Welch’s two sample t-test in Figure 7G and H.

### Data analysis of nChIP-seq

The reads from same libraries were first combined as a fastq file when they were sequenced in different lanes. We mapped the data to the human genome (hg19) using Bowtie2 with the same options as the 4C-seq (Langmead and Salzberg, 2012). Then, we generated BedGraph files for visual inspection with HOMER (Heinz et al., 2010). Peak calling was also performed with HOMER. We also mapped reads to a synthetic genomic DNA carrying the STITCH sequence inside. For this purpose, we first retrieved unmapped reads and reads that are likely to be unique from the mapped BAM file with SAMtools, with scripts of “samtools view -b -f 4” and “samtools view -b -q 10”, respectively, and combined them together, in order to remove reads that can be potentially mapped to repeat sequences. Then we re-mapped the reads against the custom reference genome. The subsequent visualization was carried out as above with HOMER (Heinz et al., 2010). We visualized the BedGraph tracks in IGV (Robinson et al., 2011). The data ranges are indicated by counts per 10 million.

---

**Figure S1.**
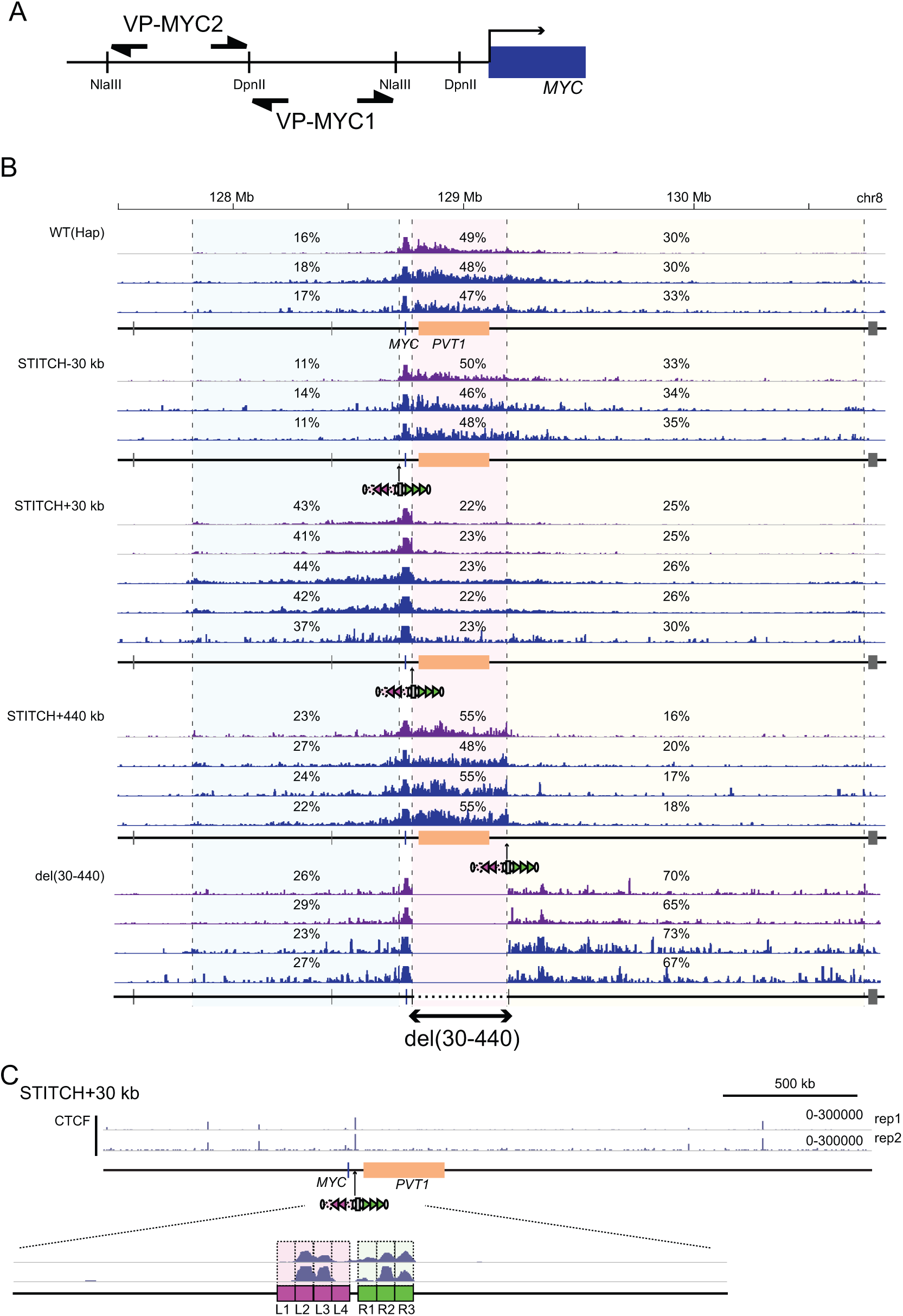
4C-seq profiles of STITCH insertion clones. (A) Schematic illustration of primer pairs (VP-MYC1 or VP-MYC2) used to amplify the 4C-seq libraries as viewpoints. (B) The 4C-seq profiles from VP-MYC1 (purple) and VP-MYC2 (blue) in different alleles. The numbers indicate the ratios of sequence reads mapped to given intervals within the locally haploid 3-Mb region around *MYC* except for the 10-kb region from the viewpoint fragment. Different tracks are results of replicate experiments. (C) nChIP-seq tracks for CTCF in the STITCH+30kb allele, mapped to a synthetic genomic DNA sequence around *MYC* with the STITCH insertion. Below is a magnified view around STITCH. The seven CTCF binding sites from L1 to R3 are marked by colored rectangles.

**Figure S2.**
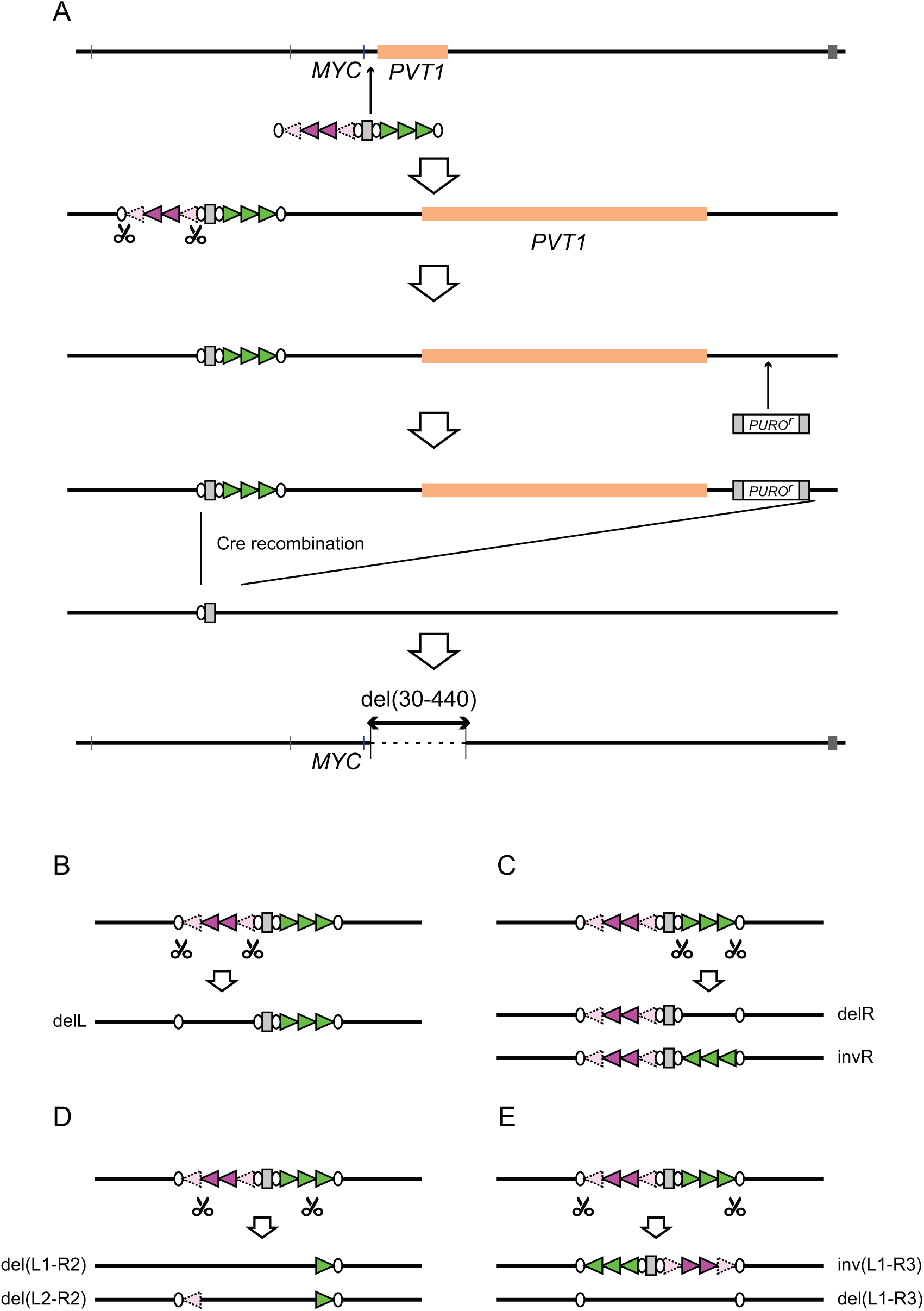
Genome editing schemes. (A) The strategy to make the del(30-440) allele. We first deleted the L1 to L4 of STITCH. Then we inserted loxP sequences together with puromycin resistant gene. Then we induced Cre expression to delete the region between the loxP sites. (B) CRISPR editing to make deletion and inversion alleles of STITCH. Guide RNAs designed against the two target sites represented by scissors were assembled with the CRISPR/Cas9 and transfected. Then we obtained the resulting clones.

**Figure S3.**
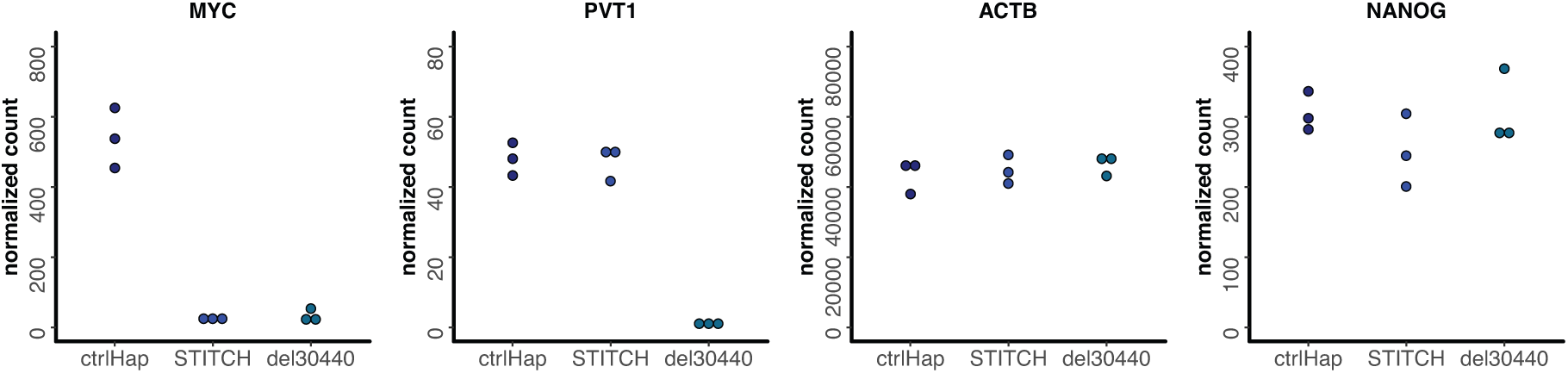
Comparison of normalized counts in RNA-seq of representative genes (*MYC*, *PVT1*, *ACTB*, *NANOG*) in Hap, STITCH+30kb and del(30-440).

**Figure S4.**
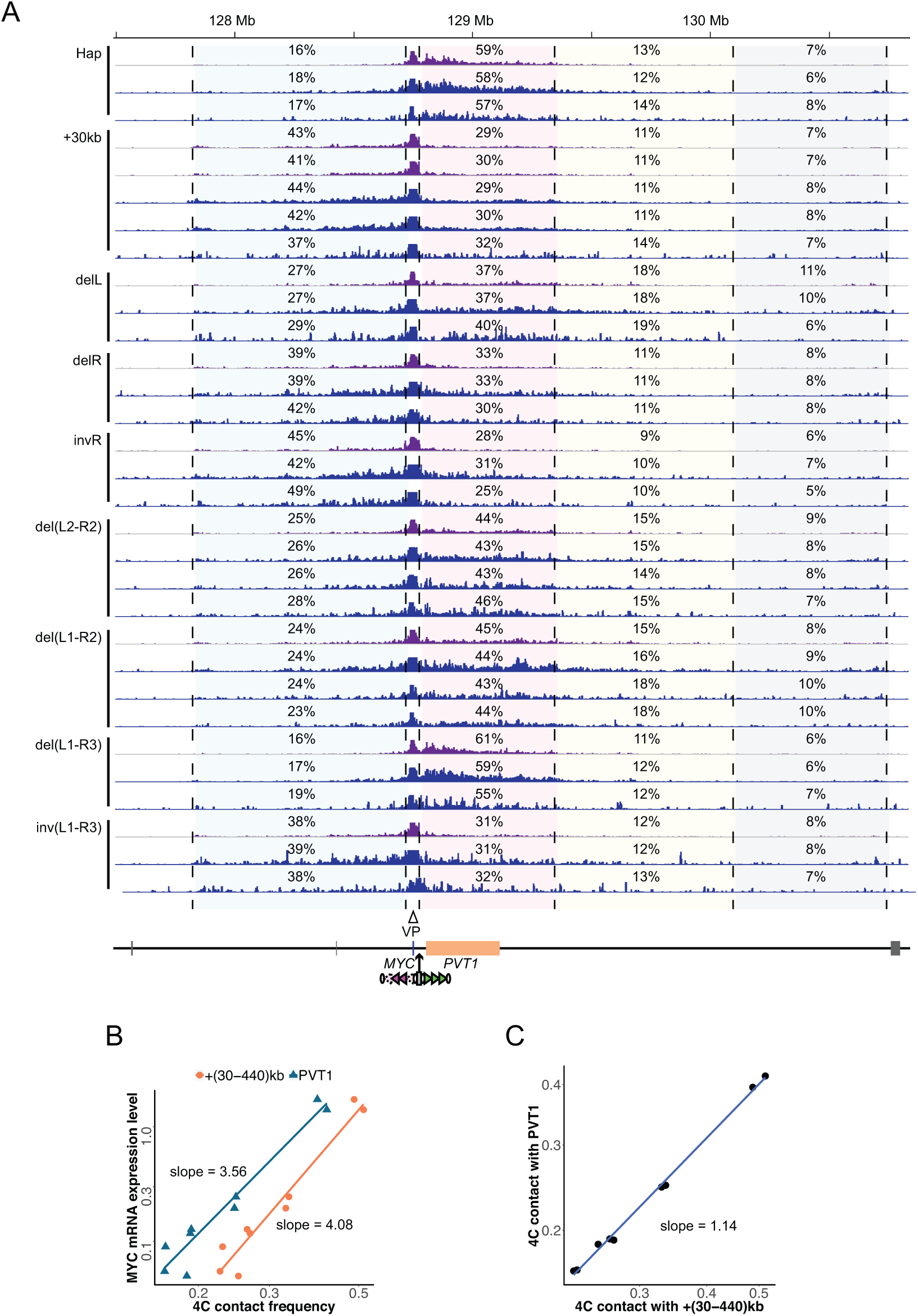
4C-seq profiles of STITCH mutants. (A) 4C-seq tracks in different alleles of STITCH mutants from VP-MYC1 (purple) and VP-MYC2 (blue). The numbers indicate ratios of mapped reads to given intervals within the locally haploid 3-Mb region except for the 10-kb region from the viewpoint fragment. (B) The log-log plot of the *MYC* expression levels against 4C contact frequency of VP-MYC1 with the +(30-440)kb region (orange) and the *PVT1* region (blue). (C) The log-log plot of 4C contact frequency of VP-MYC1 with the *PVT1* region against that with the +(30-440)kb region.

**Figure S5.**
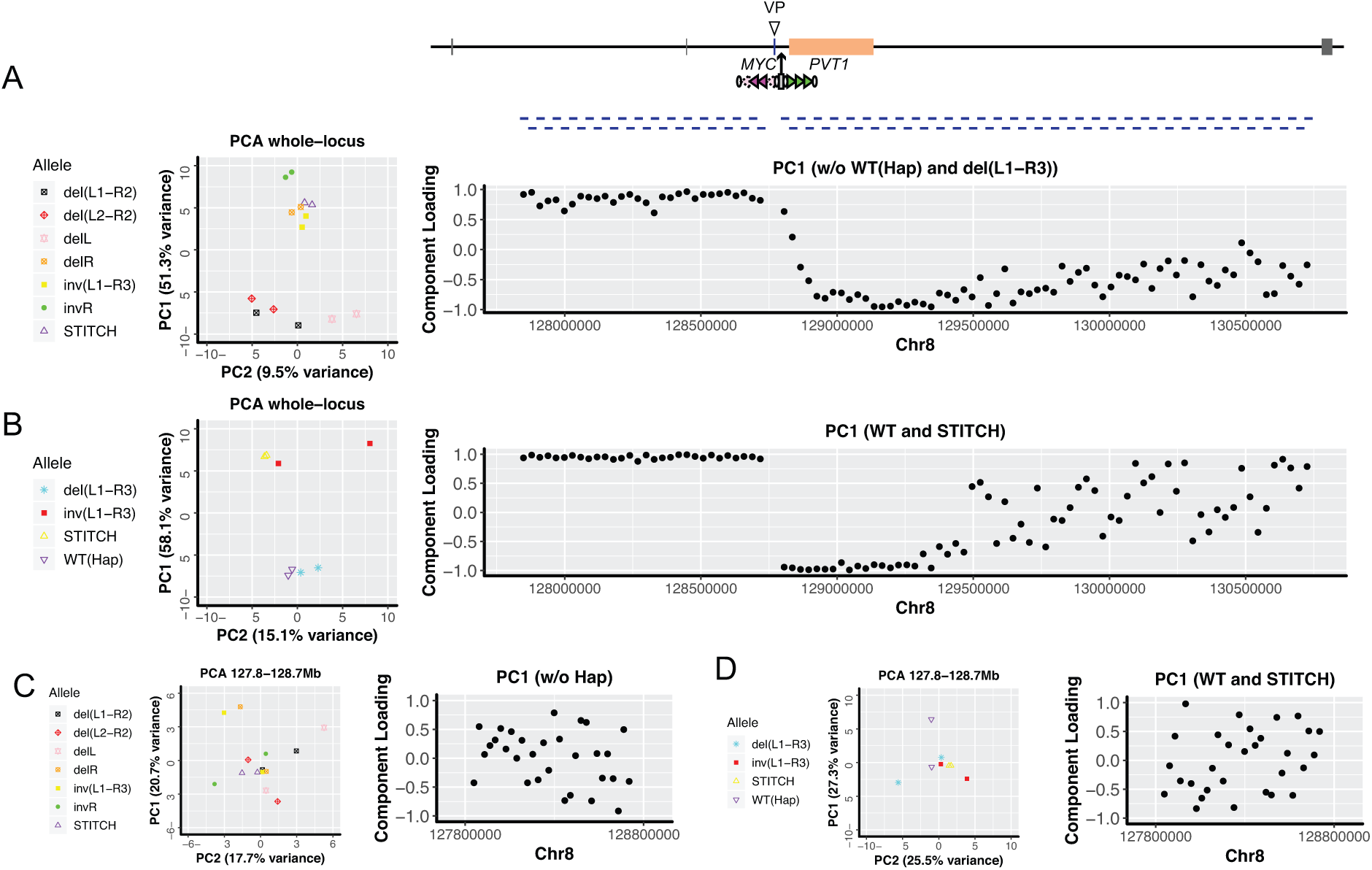
PCA of 4C-seq of STITCH mutants. (A) The PCA plot of STITCH+30kb and the mutant clones without the non-blocking alleles (Hap and del(L1-R3)) (left) and the component loadings of PC1 (right) for the whole locus. (B) The PCA plot of the non-blocking alleles (Hap and del(L1-R3)), STITCH+30kb and inv(L1-R3) (left) only, and the component loadings of PC1 (right) for the whole locus. (C) The PCA plot of STITCH+30kb and the mutant clones without the non-blocking alleles (Hap and del(L1-R3)) (left) and the component loadings of PC1 (right) for the left 900-kb region. (D) The PCA plot of the non-blocking alleles (Hap and del(L1-R3)), STITCH+30kb and inv(L1-R3) (left) and the component loadings of PC1 (right) for the left 900-kb region.

**Figure S6.**
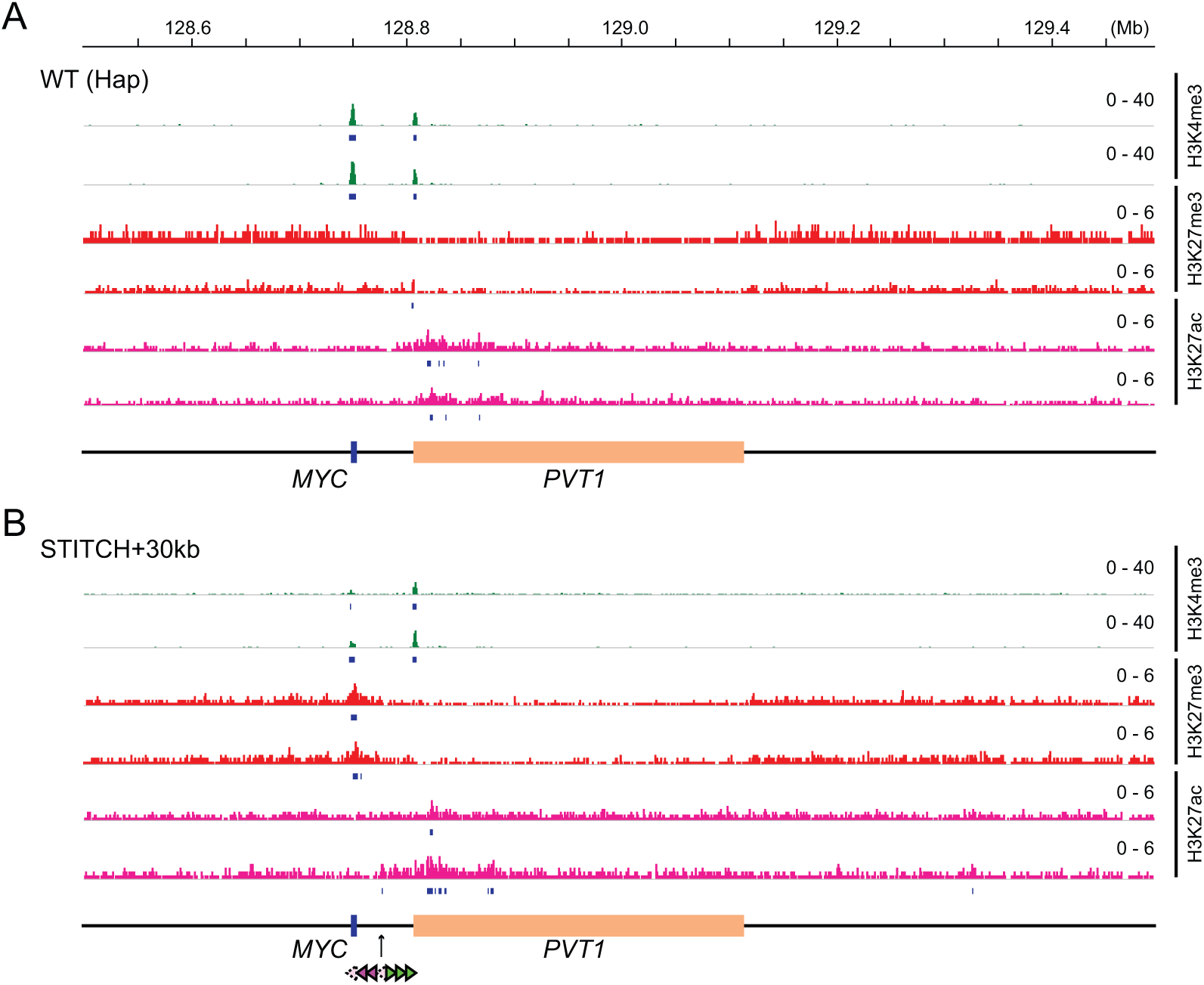
ChIP-seq profiling of Hap and STITCH+30kb alleles. (A, B) The nChIP-seq tracks for H3K4me3 (green), H3K27me3 (red) and H3K27ac (pink) of Hap (A) and STITCH+30kb (B) alleles with replicate experiments. Regions called as peaks are represented by bars below the tracks.

**Figure S7.**
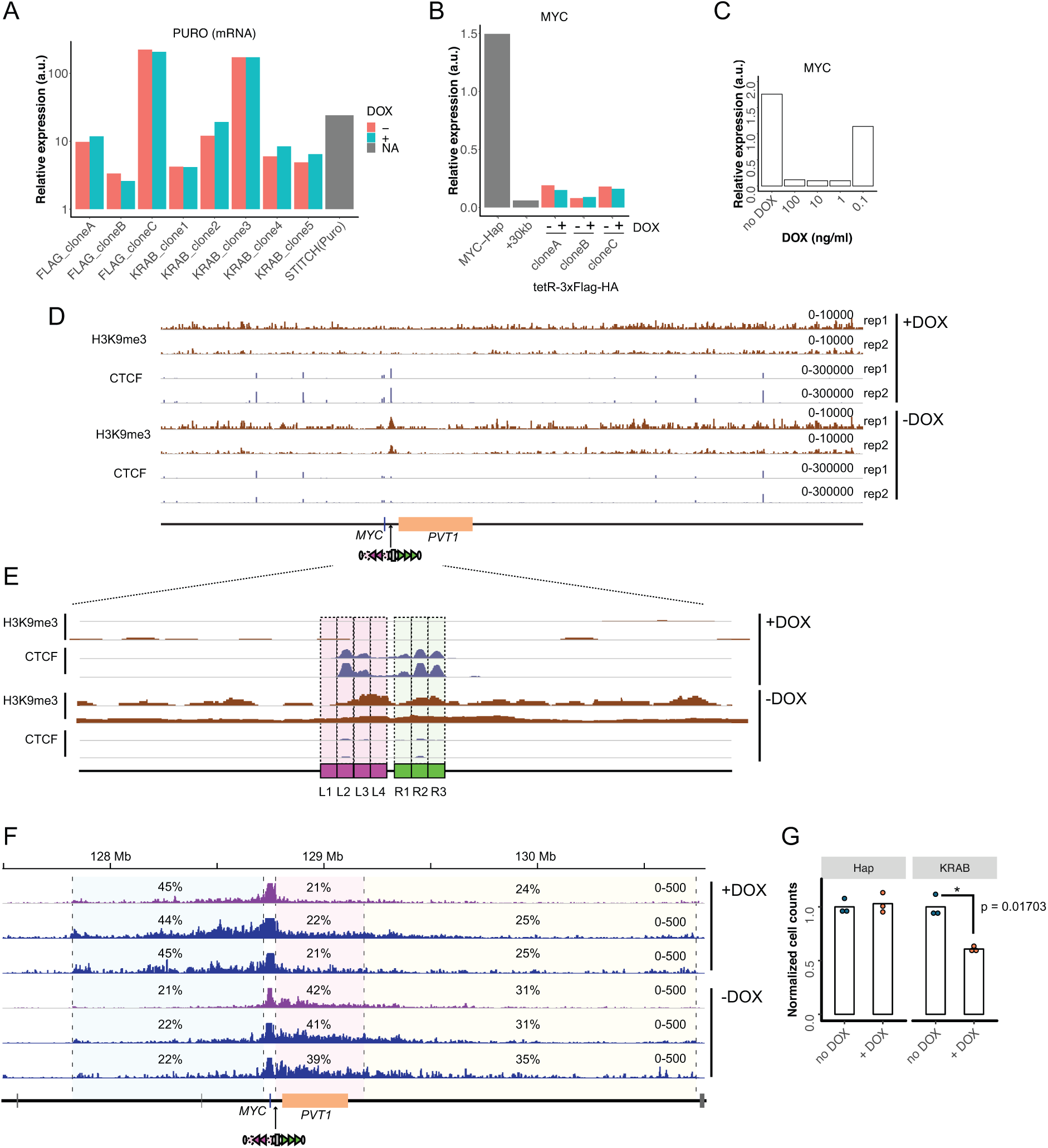
Heterochromatin induction by tetR-KRAB at STITCH. (A) The relative expression levels of the puromycin resistant gene normalized to *ACTB* in the three clones of STITCH/tetR-3xFlag-HA and the five clones of STITCH/KRAB, with and without DOX. We also compared the expression level in the STITCH+30kb clone with the selection cassette before the Cre induction. (B) The *MYC* expression levels normalized to *ACTB* in the three STITCH/tetR-3xFlag-HA clones with and without DOX. The expression levels of Hap and STITCH+30kb are the mean values in Figure 1D. (C) The relative *MYC* expression levels of the clone 3 of STITCH/KRAB with different concentrations of DOX. (D) The nChIP-seq tracks for H3K9me3 and CTCF with and without DOX. The reads were mapped to a synthetic genomic DNA sequence around the *MYC* locus with the STITCH insert. The experiments were performed in duplicates (rep1 and rep2). (E) A magnified view around STITCH of (D). (F) The 4C-seq tracks in the STITCH/KRAB clone with and without DOX from VP-MYC1 (purple) and VP-MYC2 (blue). The numbers indicate ratios of mapped reads to given intervals within the locally haploid 3-Mb region except for the 10-kb region from the viewpoint fragment. (G) Normalized cell numbers after five-days culture with and without DOX. The dots represent replicates and the bars indicate their means. * indicates p < 0.05 by Welch’s two sample t-test.

